# A Multiscale Translation of Tilman’s R* and Its Empirical Application in the Cerrado

**DOI:** 10.64898/2026.09.02.748945

**Authors:** João Augusto Alves Meira-Neto

## Abstract

Translating Tilman’s resource-ratio model (R*) to the metacommunity scale bridges local plant competition and landscape dynamics, moving ecological niche theory beyond controlled experiments. By incorporating scale dependence, stochasticity, dispersal limitation, and the stress–disturbance dichotomy, this framework also accounts for the drivers of both alpha and beta diversity. To achieve this integration, this work presents the Metacommunity Resource-Ratio Model (MetcommR) and its graphical tool, the Patchwork Biplot. This approach adapts the traditional continuous plane with a granular patch space, where each patch represents a discrete community with its area and internal variability, tracking net vectors of resource consumption and release relative to species-limiting isoclines. MetcommR articulates two fundamental regimes. In disturbance-governed metacommunities, biomass loss releases resources, interrupting depletion and preventing competitive exclusion, which promotes coexistence via the competition– colonization trade-off and maintains alpha and beta diversity. Conversely, in stress-governed systems, biomass accumulation and severe resource scarcity push communities toward isoclines, favoring the monodominance of tolerant species through competitive exclusion under the tolerance–fecundity trade-off. We demonstrate the model’s empirical utility through a Cerrado case study in the Paraopeba Reserve. Analyzing canopy openness (light) and soil nitrogen reveals that long-term fire suppression induces light stress, elevating competitive exclusion risks for shade-intolerant species. Consequently, the Patchwork Biplot serves as a tool for evaluating ecological dynamics and vulnerabilities to guide management and conservation strategies.

## Introduction

### The MetcommR model

Integrating resource competition models with the demands of modern landscape ecology is an increasing priority. Developed by David Tilman building on niche theory in his foundational works (Tilman, 1986, 1988), the R* resource competition model established a widely accepted mechanistic foundation for understanding interactions among resource-consuming competitors and their direct impacts on the environment. The R* value represents the minimum level to which a monoculture species can deplete a limiting resource. When competition involves two limiting resources, the competitive outcome is determined by the intersection and positioning of the species’ zero-net-growth isoclines, together with the resource consumption and supply vectors in a two-dimensional space (Tilman, 1988). Thus, functional and physiological plant traits directly influence competitive outcomes through their role in reducing environmental resource levels. In Tilman’s original formulation (Tilman, 1988), competition is modeled assuming that resource ratios (R1 and R2) locate a community at a single point within a perfectly continuous, smooth two-dimensional space. The model operates strictly at the local community scale, where direct plant-to-plant competitive interactions occur. At this local sampling scale—typically on the order of square meters or experimental plots—the disturbance regime and initial resource availability are assumed to be fixed and constant for the community under analysis.

Although the classic R* model serves as a mechanistic foundation for interspecific competition, the original theory lacked a hierarchical spatial structure and graphical-statistical tools capable of representing metacommunities and analyzing multiple spatial grains (local communities) simultaneously across scales (Leibold et al., 2004). This gap between local models and cross-scale complexity exemplifies what Tilman (Tilman, 2007) highlighted as essential for advancing ecological theory: *“Plant ecology needs closer links between analytical theory, observations and experiments. Simple verbal theories can generate novel ideas but the logical implications of such scenarios are best explored using the rigorous logic of mathematics. Predictions of theory can then be tested via experiments and comparative studies.”* Using the R* model as a foundation, expanding to a multiscale approach helps to fulfill this vision. By incorporating scale dependence, resource stochasticity, and spatial dynamics across multiple environmental planes, the Patchwork Biplot connects the mathematical logic of the original R* model to empirical observations across scales, from local plant-to-plant interactions to the landscape-level dynamics of MetcommR. Consequently, both frameworks share the same fundamental design (Figure 1).

**Figure 1.**
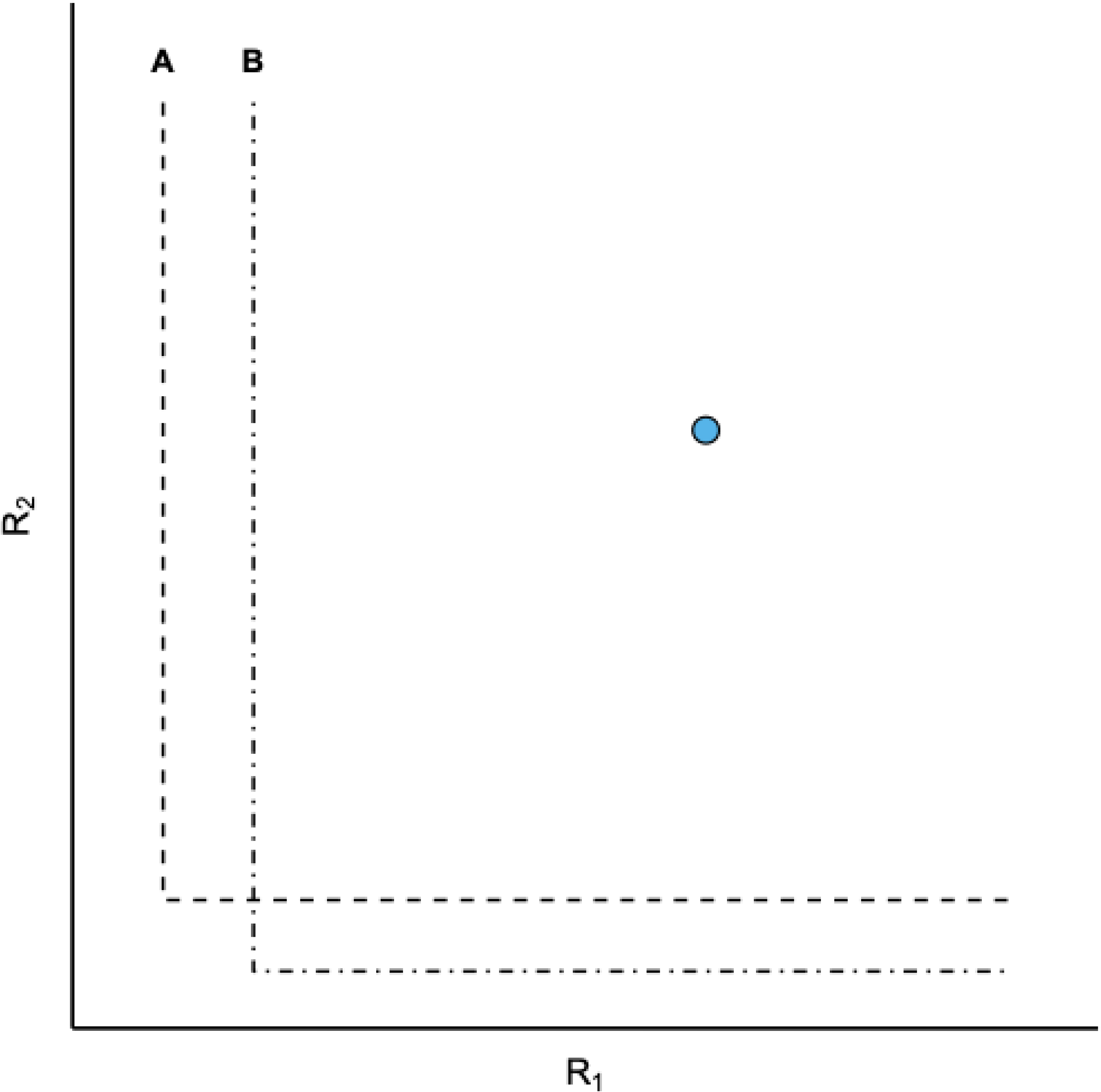
Both R* (Tilman 1988) and MetcommR start from the same fundamental design. The point (R*) or the circle (MetcommR) has its proportion of R1 and R2. The isoclines, in both cases, are determined by prior information on the limitations of these species in other communities (R*) or metacommunities (MetcommR).

The R* model builds upon Hutchinson’s niche concept (Hutchinson, 1957), whose simplest formulation is a two-dimensional space defined by tolerance limits along two orthogonal environmental axes. Tilman’s resource model (Tilman, 1988) adopts this deterministic niche structure, predicting competition in a two-species community where resource ratios (R1 and R2) locate the community at a single point within a smooth, two-dimensional plane. Although plant development requires numerous resources, both Tilman’s model and the proposed Patchwork Biplot evaluate resource consumption and release-pairwise focusing on a single community within a metacommunity, a pair of species, and a pair of resources in their foundational configuration (Figure 1). Both frameworks use this two-species baseline to establish core principles before expanding into multispecies extensions. Modeling more than two resources requires multidimensional spaces, analogous to Principal Component Analysis (PCA) ordination plots, which are displayed as pairwise projections to facilitate interpretation.

Environmental resource availability exhibits self-similarity across spatial scales in a granular, patchy pattern. This availability depends on the scale of observation, as interacting edaphic and topographic factors influence nutrient and water retention and flux across spatial extents (Kravchenko et al., 2000; Biswas, 2019). At the plant-to-plant scale (meters), micro-topography, bioturbation, and organic matter condition fine-particle fractions to regulate immediate water, nitrogen, and phosphorus retention. At intermediate scales (hectares), slope position (crest, mid-slope, footslope) and erosional processes determine water dynamics via soil clay content. At the landscape scale (square kilometers), geomorphic landforms and parent material properties govern water availability through variations in coarser fractions like silt and sand (Kravchenko et al., 2000; Zeleke & Si, 2006).

Consequently, the primary drivers of soil texture and moisture vary with the sampling scale (e.g., 1, 100, or 10,000 m), causing distinct plant life forms and communities to respond to scale-specific edaphic gradients. For instance, a meter-scale model can be parametrized for Therophytes, Geophytes, or Chamaephytes (Raunkiaer, 1934) plant communities, whereas a forest plot sampled for trees (phanerophytes) requires a distinct, scale-appropriate model. Each model possesses its own spatial granularity (communities) determined by its focal scale within the metacommunity. By replacing a single continuous plane with a granular, scale-dependent patch space, the extended Patchwork Biplot model defines each “patch” as a discrete community at the chosen scale. This establishes the mechanistic bridge required to connect species coexistence across spatial scales, from local plant-to-plant interactions up to multi-metacommunity dynamics.

In the Patchwork Biplot, a species’ isoclines are defined by the lowest R1 and R2 resource concentrations observed across all communities in which that species occurs within the focal metacommunity (Figure 2). This empirical determination differs fundamentally from the zero-net-growth isoclines of Tilman’s R* model (Tilman, 1988). Here, the focal metacommunity—or a combined set when comparing multiple metacommunities—represents the universe of analysis. Consequently, the minimum R1 value recorded among all communities harboring the species defines its R1 isocline boundary. Repeating this procedure for R2 establishes the corresponding R2 boundary, with their intersection marking the vertex of the species’ isoclines in that metacommunity. Thus, isocline determination in the Patchwork Biplot contrasts with the experimental measurement of R* isoclines, which require establishing an empirical balance between growth and loss (or disturbance) at specific resource levels. Although consumption and release vectors in the Patchwork Biplot can be assumed to operate near equilibrium, this balance is inferred rather than directly measured. Furthermore, while the model assumes a baseline level of intrinsic loss associated with routine plant phenology (e.g., leaf shedding, fruit drop, seed dispersal, or bark sloughing), these background dynamics do not define the isoclines. Similarly, environmental fluctuations and altered disturbance regimes that would shift isocline positions in the classic R* model do not alter isoclines in the Patchwork Biplot; instead, they shift the resource vectors. While both frameworks can portray multiple communities within a metacommunity regardless of species composition, abundance, or biomass, only MetcommR can simultaneously evaluate multiple metacommunities. Finally, whereas communities in Tilman’s R* model are depicted as dimensionless points, they are represented as circles with defined spatial properties in the Patchwork Biplot.

**Figure 2.**
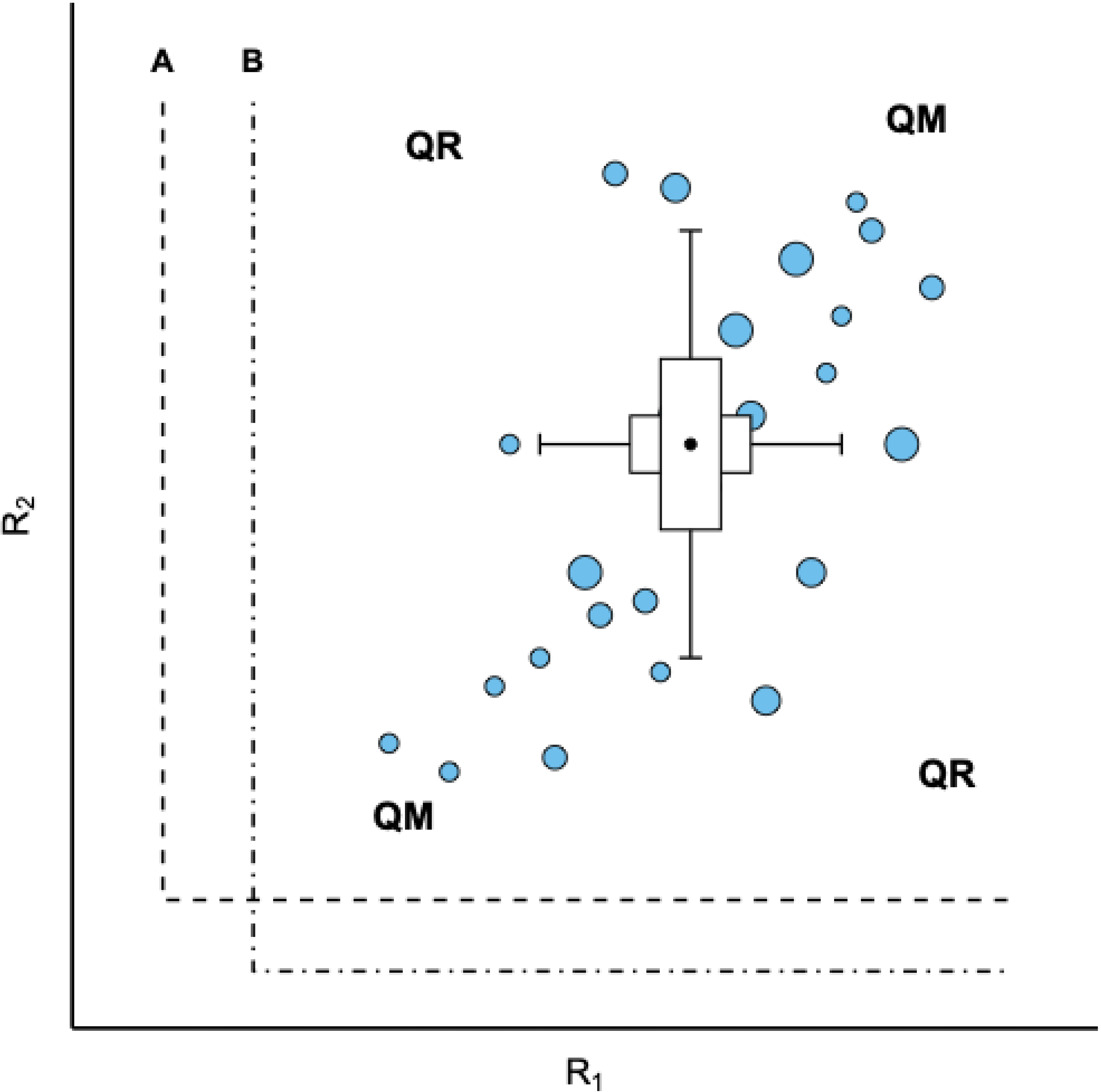
The model of the Patchwork Biplot and its parts. In its simplest version, we have a biplot of two resources, R1 and R2, with communities of different areas (represented by the sizes of the circles), in this case a metacommunity governed by the disturbance, with communities far from the limiting values of the isoclines. Two species define the isoclines according to known minimum values of their preferences in this or other metacommunities for which data are available. Therefore, we can insert data from one or more metacommunities for analysis. The mainstream quadrants (QM) show the diagonal of the metacommunity dynamics when resources R1 and R2 are consumed or released simultaneously. The rare quadrants (QR) show the diagonal of the metacommunity dynamics when resource R1 is consumed while resource R2 is released, and vice versa.

Circle areas correspond to the spatial footprint (e.g., m^2^, hectares, km^2^) of each community within the metacommunity, serving as a proxy for its potential internal variability at finer scales. Larger circles indicate a greater potential for internal heterogeneity in R1 and R2 when analyzed at downscaled levels. The Patchwork Biplot accounts for resource proportions in each community at a specified scale, enabling statistical evaluation of the metacommunity based on community positions along the R1 and R2 axes. This variability is summarized by two orthogonal boxplots: the horizontal boxplot describes the distribution of R1 across communities, while the vertical boxplot depicts the distribution of R2. Intersecting at their medians, these boxplots divide the granular two-dimensional space into four quadrants. Each patch (circle) represents a discrete community composed of one or more species, while the boxplots display the medians and quantiles of the overall metacommunity. Figure 2 illustrates this framework for a representative two-species metacommunity.

As Leibold et al., (2017) emphasized, adopting a landscape perspective on the role of biota in community assembly and ecosystem processes is essential for advancing ecological understanding. The MetcommR aims to support this perspective. In the Patchwork Biplot, Mainstream Quadrants (QM) define vector trajectories when both resources are consumed or released simultaneously. The off-diagonal Rare Quadrants (QR) represent scenarios where one resource is consumed while the other is released. When metacommunity dynamics traverse the Mainstream Quadrants (QM), the absence of disturbance drives communities toward the lower-left corner as plant growth depletes both resources. Conversely, disturbance shifts communities toward the upper-right corner as biomass mortality and decomposition increase resource availability. When dynamics traverse the Rare Quadrants (QR), one resource is consumed while the other is released, driving trajectories toward the upper-left or lower-right corners. This behavior corresponds to systems where disturbance involves net resource export, such as livestock grazing with biomass removal, nutrient volatilization by fire, or timber harvesting.

### Metacommunities ruled by disturbance

Within communities, deaths (or partial biomass loss) generate vectors opposite to growth vectors because biomass decay releases resources. While random individual mortality has negligible overall effects, mortality driven by widespread disturbance profoundly impacts the model by amplifying vector magnitudes or reversing dynamic resource vectors across communities and metacommunities.

The model can be extended to multispecies scenarios, incorporating as many isoclines as there are species (e.g., species A, B, and C in this metacommunity example). Each community (grain) occupies a distinct position in the biplot, contributing to the overall metacommunity’s quantiles, confidence intervals, and outliers. Although community spatial area is not currently weighted in vector or boxplot calculations, incorporating area-weighting represents a promising future expansion.

In multispecies models, immigration acts as a key process counteracting dispersal limitation as metacommunities and expanded niche models (Leibold et al., 2004; Soberón, 2007). Higher immigration rates increase compositional similarity among communities (Hubbell, 2001), producing more parallel resource consumption vectors. Conversely, as community compositions and abundances deviate from the metacommunity mean, differential consumption ratios of R1 and R2 become more likely, causing resource vectors to diverge from parallelism.

A shift in vector orientation toward the nearest limiting isocline can favor one species while driving another to local exclusion. Consequently, each grain exhibits unique dynamics and distinct probabilities of dominance by species A, B, or C (Figure 3). Furthermore, disturbance regimes can fluctuate independently within each community. The model reveals that R1 and R2 consumption and release during plant growth are tightly linked and proportional, driven by the consumption ratios of dominant species across the metacommunity. Under high immigration rates, individual community vectors move nearly parallel to the metacommunity mean vector. Conversely, under low immigration rates, distinct dominant species across communities generate resource vectors that diverge from the mean.

**Figure 3.**
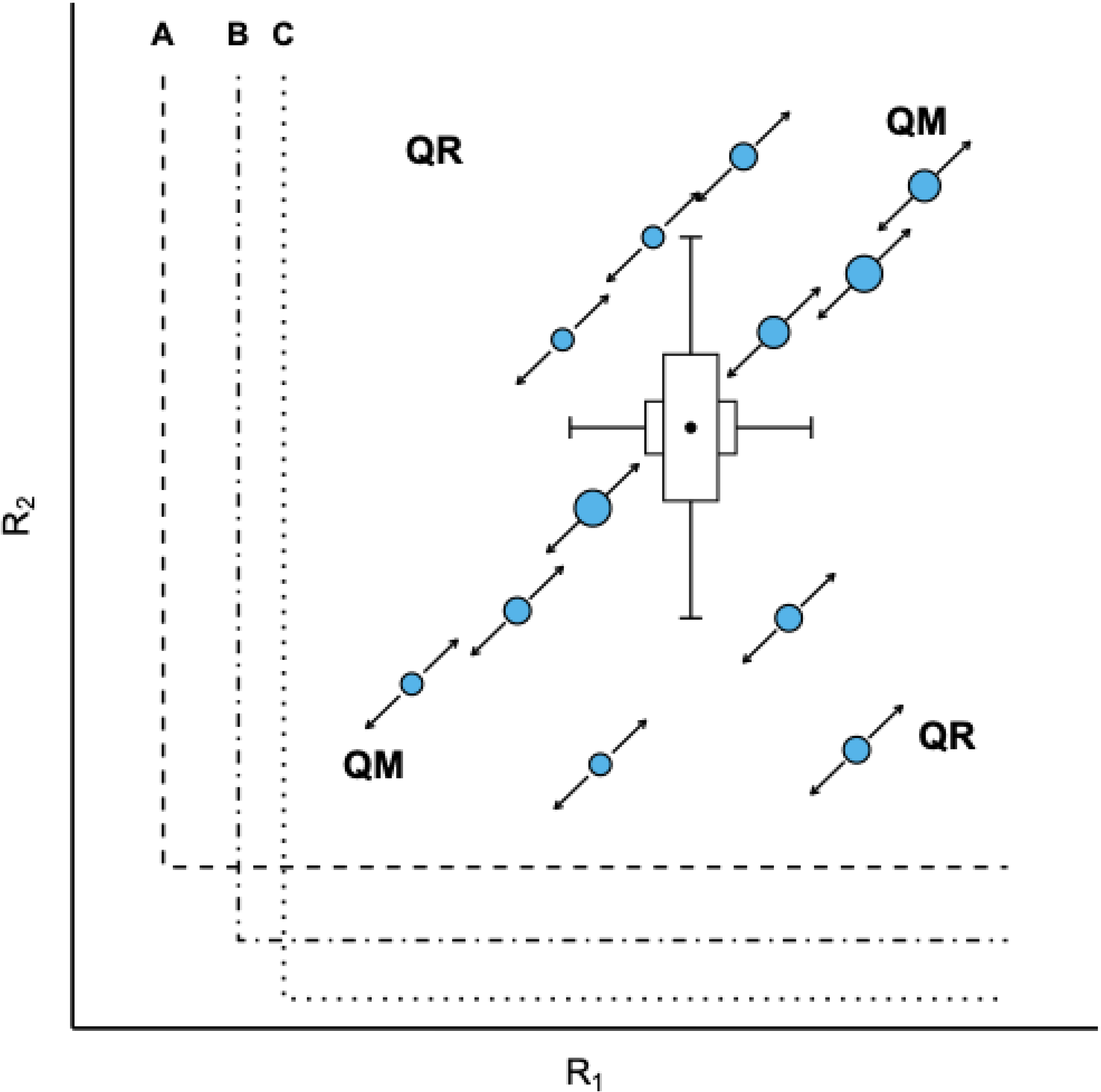
Patchwork Biplot of a disturbance-ruled metacommunity with two resources and three species. The two resources are consumed or released concomitantly. The metacommunity dynamics are determined by the diagonal of the mainstream quadrants (QM). This example is of a metacommunity highly connected by immigration, without isolated communities. High immigration confers high predictability of community dynamics due to the very similar relative consumption of R1 and R2 provided by the very similar species compositions and abundances, generating parallel vectors. The diagonal of the Rare Quadrants (QR) does not define the dynamics in this case.

The net balance between resource consumption and release, driven by species composition (A, B, and C), generates the community vectors that rule metacommunity dynamics: a net consumption vector pointing toward the lower-left corner of the model, or a net release vector pointing in the opposite direction.

When one resource is depleted while another is released, and vice-versa, metacommunity trajectories shift toward the Rare Quadrants. Under these conditions, only one resource induces stress at time, R1 or R2 (Figure 4). Nevertheless, continued depletion of a single limiting resource will eventually drive the metacommunity into a state of generalized stress, resulting in deterministic monodominance by a single species, as in the Figure 4. When a system is ruled by the stress inherent in progressive biomass accumulation and resource scarcity (e.g., intense light limitation in forest understories), communities are driven toward limiting isoclines. Under such environmental constraints, resource scarcity can promote competitive exclusion, stabilizing communities into monodominance by highly stress-tolerant species.

**Figure 4.**
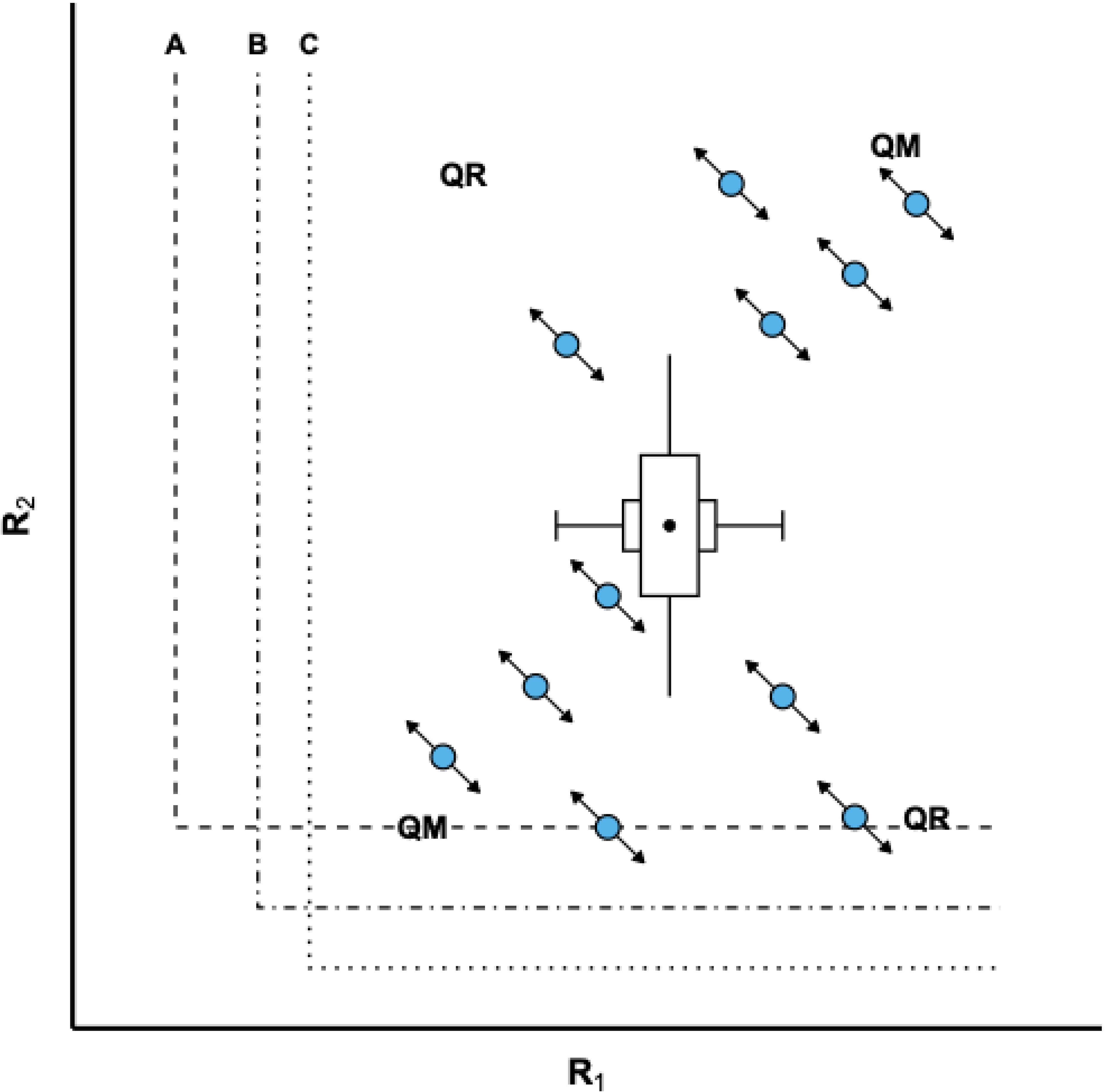
Patchwork Biplot exemplifying a situation where the metacommunity dynamics are determined by the consumption of R2 while R1 is released. The vectors are on the diagonal of the rare quadrants (QR). The dynamics of this community will lead to future governance through the stress of R2. Only disturbance can alter the dynamics, but in the opposite direction. In this case, the diagonal of the mainstream quadrants (QM) does not define the metacommunity dynamics.

In Figure 4, vector shifts occur as one resource is consumed while the other is released. A community vector shifting toward the upper-left quadrant favors species A, which better tolerates low levels of R1. Conversely, a shift toward the lower-right quadrant favors species C, which tolerates lower levels of R2. However, each patch (grain) exhibits distinct dynamics and varying probabilities of dominance by species A, B or C. If a widespread disturbance steers all community trajectories toward conditions favoring species A, species A can achieve monodominance across the metacommunity. This represents an extreme case of competitive exclusion at the metacommunity scale, where no mechanism for stable coexistence is predicted.

In resource space, vector dynamics are governed by the net balance between resources consumption and release across communities. When mortality and biomass loss occur, the resulting release of resources generates community vectors pointing away from limiting isoclines. By steering the system away from severe limitation, these vectors prevent competitive exclusion and foster opportunities for coexistence.

In the Patchwork Biplot, species-specific consumption rates can be specified for each community to determine the orientation of its trajectory within two-dimensional resource space. Variations in species composition, relative abundance, and biomass enable each community to exhibit distinct vector dynamics. Under high immigration rates, consumption and release dynamics become increasingly homogeneous across the metacommunity because unrestricted dispersal aligns local species compositions and relative abundances. Consequently, the continuous influx of propagules minimizes compositional and abundance dissimilarity between local communities and the regional pool. As a result, proportional resource (R1 and R2) uptake and release by regionally dominant species cause individual community consumption vectors to move roughly parallel to one another.

In contrast, when dispersal is limited and immigration rates are low, neutral drift can become an important driver (Hubbell, 2001). In these scenarios, stochastic fluctuations can lead to local monodominance by distinct species across different patches (grains). Because each species exhibits intrinsic, species-specific proportions of R1 and R2 consumption and release, this local monodominance generates divergent resource vectors that do not align with those of other communities. By driving dynamics toward peripheral positions in the biplot, such disparate trajectories establish unique resource ratios that favor specific niches and distinct isocline intersections (“corners”), ultimately supporting the persistence of rare species across the metacommunity (Figure 5).

**Figure 5.**
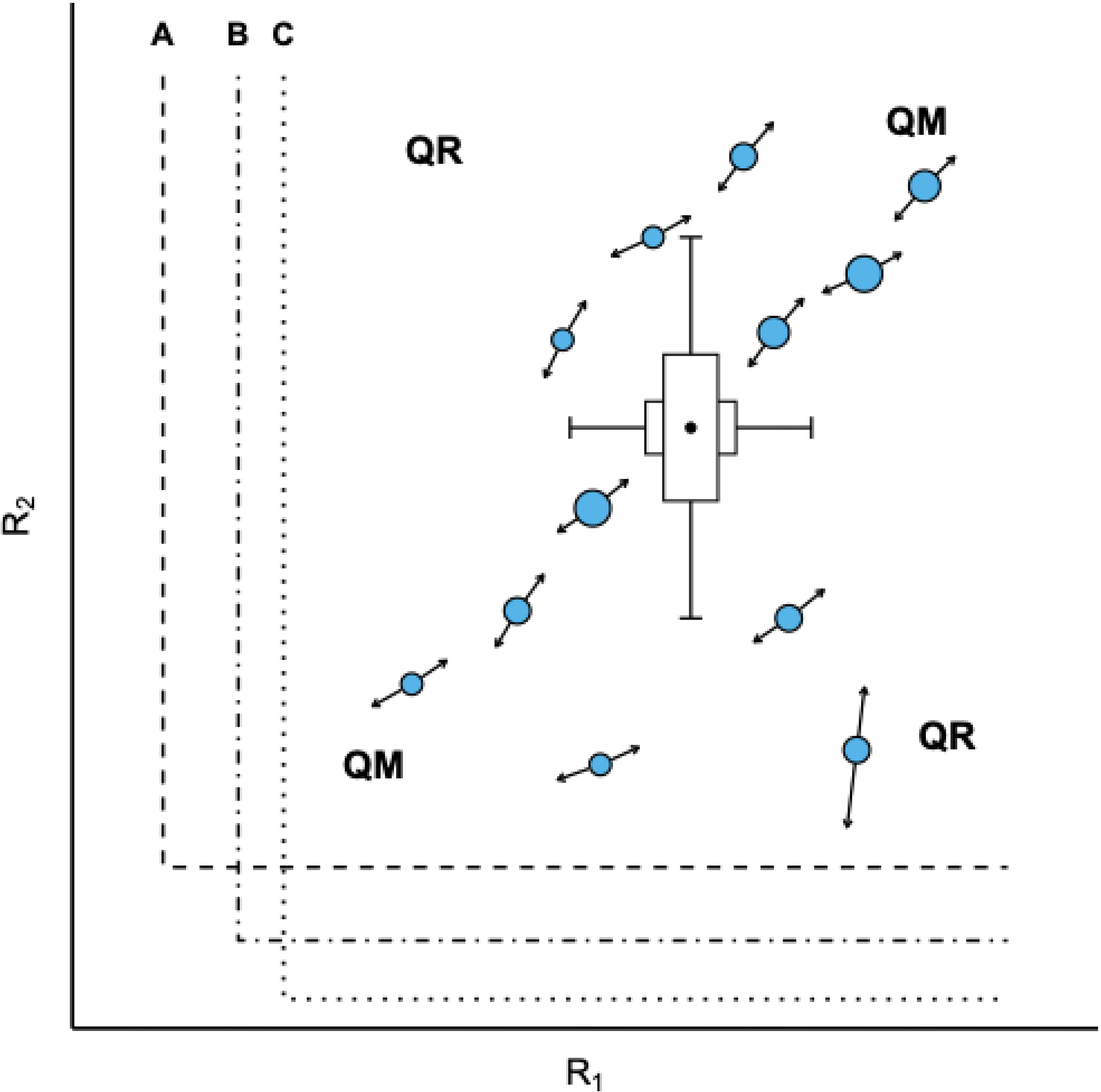
Patchwork Biplot exemplifying a metacommunity with severe dispersal limitation, with simultaneous consumption and release of resources. The low immigration rate tends to make communities monodominant due to drift neutrality (Hubbell, 2001). Different species may consume R1 and R2 resources in different proportions, and monodominant communities within a metacommunity have less predictable dynamics. Their vectors are less parallel than in communities with a high immigration rate.

In an isolated community, a high relative abundance of species A can generate a consumption and release trajectory that diverges significantly from the rest of the metacommunity. Although a higher dominance of species A could also be driven by deterministic processes (e.g., density-dependent effects), it is used here to illustrate neutral dynamics resulting from low immigration (Figure 5).

Another consequence of dispersal limitation is that a species’ isoclines may be located far from points of extreme resource depletion, simply because the species does not occur where the resource approaches its limiting threshold.

In a metacommunity with high species richness and high immigration, the neutral framework described by Hubbell (Hubbell, 2001) predicts that local dominant species will match regionally dominant species. Conversely, under strong dispersal limitation (low immigration), local dominants frequently diverge from regional dominants as a result of neutral drift. This generates diverse localized conditions that help explain the persistence of numerous rare species across the metacommunity, particularly if immigration is subsequently re-established. Consequently, even regionally dominant species can face local competitive exclusion when a community transitions from a disturbance-governed to a stress-governed regime (Figure 5).

An isolated local community exhibiting incomplete monodominance of species A due to neutral drift consumes R2 at a higher rate relative to R1 (or vice-versa) than the metacommunity average. Consequently, its resource vector diverges from those of neighboring communities, driving the system toward the limiting isocline of species A before reaching the isoclines of species B or C. Under complete monodominance of A species, the community intersects species A’s isocline and stabilizes, oscillating along the boundary as biomass growth and disturbance alternate.

Under incomplete monodominance, sparse individuals of species B and C gain a competitive advantage by drawing R2 concentrations below species A’s isocline. This drives mortality in species A via competitive exclusion, overriding the initial dominance established by neutral drift. As the community transitions from a disturbance-governed to a stress-governed regime, deterministic competitive dynamics displace species A, shifting dominance toward species B and ultimately species C. During this transition, the community’s resource trajectory oscillates along species A’s isocline, where mortality in species A releases resources (upward vector) while the growth of species B consumes resources (downward vector), while sliding toward species B’s R1 isocline as species A declines. Whether deterministic monodominance by species B or C becomes complete depends on whether R1 is depleted to species B’s or C’s isocline threshold prior to species A’s local extinction. Thus, in a multispecies biplot lacking immigration, neutral drift (Hubbell, 2001) can generate distinct resource consumption trajectories, producing localized monodominances characterized by species-specific R1 and R2 consumption and release ratios.

### Metacommunities ruled by disturbance

In disturbance-governed systems, localized or widespread biomass loss releases resources and shifts communities away from limiting isoclines, alleviating continuous resource depletion pressure. In this disturbance-dominated regime, the Competition–Colonization Trade-off (Cadotte, 2007), a trade-off ruled by disturbance, can predominate wherein intermediate disturbance frequencies or intensities maximize alpha and beta diversity by structuring the landscape as a mosaic of patches (communities) with varying biomass levels. Intermediate disturbances prevent monodominance by competitively superior species (low-disturbance specialists) that require temporal stability to monopolize resources. Concurrently, disturbances continuously create colonization opportunities essential for maintaining fast-growing pioneer species with high dispersal capacity (high-disturbance specialists). Thus, the equilibrium between localized mortality and recolonization sustains the coexistence of species operating at opposite ends of this functional trade-off, fully aligning with the predictions of the Patchwork Biplot Model.

In a metacommunity experiencing oscillating disturbance regimes within the shared tolerance zone of species A, B, and C (away from isoclines and therefore not ruled by stress), immigration rates determine the alignment of community trajectories. High immigration aligns local species compositions and abundance proportions with metacommunity averages, prompting individual community vectors to run parallel to the metacommunity mean vector. Conversely, low immigration rates result in divergent, non-parallel trajectories across communities. Intermediate disturbance and high immigration rates interact synergistically within disturbance-ruled metacommunities. Intermediate disturbance enhances species richness (alpha-diversity) across local community and metacommunity scales, as spatial variation in disturbance intensity facilitates species coexistence (Cadotte, 2007). High immigration rates continually supply local communities with species and individuals from the metacommunity pool (Hubbell, 2001), thereby enhancing alpha-diversity.

Consider a single community undergoing succession under a low-disturbance regime. Early in succession (1), high availability of R1 and R2 combined with rapid resource uptake produces a large vector directed toward the coexistence corner (where the isoclines of species A and B intersect). Over time (2, 3), this consumption vector shrinks as the system approaches a stress-governed state, ultimately reaching equilibrium (4), where resource uptake balances total disturbance (including intrinsic phenological loss). Here, the boxplots depict the distribution of positions occupied by this single community across successive time steps. Conversely, under a scenario of intensifying disturbance initiated from position 4 (e.g., driven by global change), the trajectory reverses from position 4 back to position 1. Under these conditions, resource release vectors are largest at position 4 and progressively diminish toward position 1 (Figure 6).

**Figure 6.**
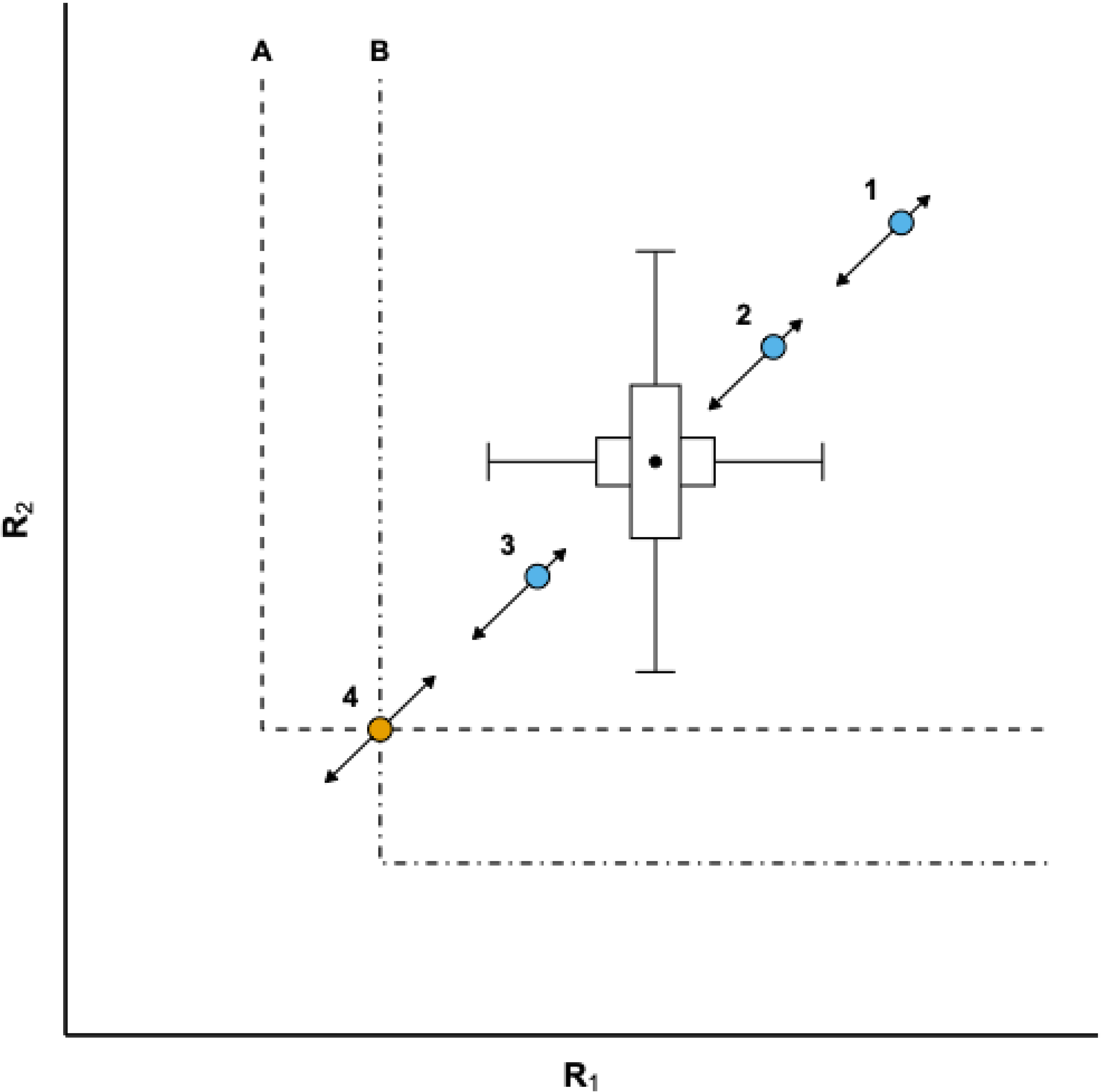
In a Patchwork Biplot, a metacommunity can depict the same community at several different times. The example shows four distinct successional moments in the same forest community where both the consumption and release of resources R1 and R2 occur simultaneously. At time 1, the community is governed by intense disturbances that provide greater availability through the release of R1 and R2, with the resource consumption vector increased, leading the community to a rapid accumulation of biomass. At times 2 and 3, the decrease in resources allows for biomass accumulation in a dynamic that leads the community to stability as it becomes governed by the stress of both resources, at the intersection of the isoclines A and B for R1 and R2.

Within the Patchwork Biplot model, biodiversity regulation operates across local and landscape scales. Alpha diversity is maintained locally by disturbances that prevent competitive exclusion, promoting plant-to-plant coexistence at the microscale mediated by functional trade-offs. High immigration rate also connects the metacommunity species pool to the local species compositions, improving alpha-diversity (Hubbell, 2001). Conversely, beta diversity is generated by edaphic spatial heterogeneity, reflected in the distinct positioning of patches (grains) in the biplot, that enable different paths that, even when parallel, wind up in different points of the isoclines. This deterministic path, combined with dispersal limitation and high (or low) disturbance regimes across the landscape, integrates coexistence mechanisms with spatial environmental segregation to explain variations in community composition, alpha-diversity and metacommunity betadiversity.

### Stress-Ruled Communities

In forest metacommunities governed by stress, where biomass accumulation causes severe light limitation in dense understories, community structuring aligns with the Tolerance–Fecundity Trade-Off (Muller-Landau, 2010). This trade-off predicts that under severe light limitation, large-seeded species allocate greater energy and nutrient reserves per propagule, conferring high tolerance to low-light stress beneath closed canopies at the expense of numerical seed yield (fecundity). In contrast, small-seeded species prioritize high fecundity and widespread dispersal, efficiently colonizing microsites where temporary light gaps occur. Thus, fine-scale stress heterogeneity within community patches (at the microsite scale) fosters tolerance–fecundity coexistence and reduces the likelihood of local competitive exclusion. Consequently, the absolute competitive exclusion observed experimentally by Tilman (Tilman, 1988) becomes scale-dependent, marking a fundamental distinction between the classic R* model and the Patchwork Biplot framework.

Although disturbances still occur, stress remains the primary driver in these communities, rendering consumption and release vectors negligible. In Figure 7, upon reaching the tolerance boundary on species B’s isocline (limited by low R2 availability), disturbance-driven oscillations in R2 cause the community to slide along the isocline toward progressive R1 depletion (Trajectory 1). This trajectory occurs in open ecosystems with net R1 loss or where the two resources operate independently. Conversely, if R1 enters the system from external inputs, the trajectory reverses toward R1 accumulation (Trajectory 2) (Figure 7).

**Figure 7.**
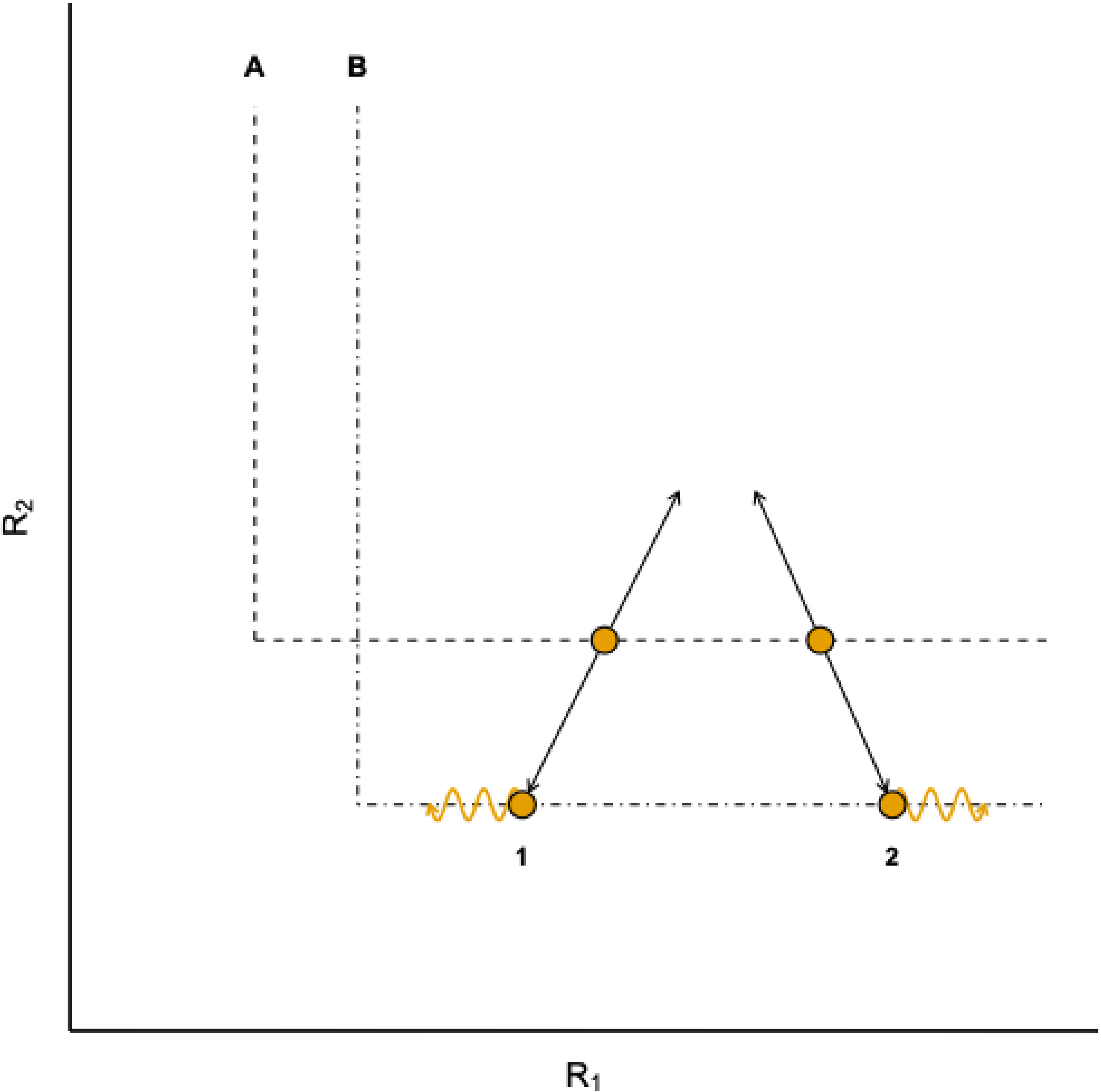
A detail of the Patchwork Biplot. An isolated community in a Patchwork Biplot exemplifying its dynamics on the isocline of species B, which is more tolerant to the stress of resource R2. 1) With a dynamic of consuming both resources simultaneously, the dynamics through the mainstream quadrants tend to slide the community to the left along the R2 isocline of species B. 2) With a dynamic of consuming one resource while another is released, and vice versa, the dynamics through the rare quadrants tend to drift the community sliding to the right along the R2 isocline of species B.

Under reduced disturbance, stress-governed communities stabilize into complete monodominance (with negligible oscillations) along limiting isoclines, regardless of immigration rates or neutral drift. Although the Patchwork Biplot Model contextualizes the absolute competitive exclusion predicted by Tilman (Tilman, 1988), monodominance can still emerge under neutral drift (Hubbell, 2001). However, in deterministic cases, the absence of relevant disturbance could reduce local diversity to the single most tolerant species of each community. Consequently, communities governed solely by stress tend to lose alpha diversity, though beta diversity may persist across the broader metacommunity.

Under predominant stress and biomass accumulation, dynamic net vectors vanish because net growth is nearly zero and biomass losses tend to be immediately replaced, maintaining overall biomass nearly constant. Only severe, widespread disturbance can disrupt this stability by releasing R1 and R2 resources. Thus, in a metacommunity historically governed by stress, disruptive disturbance serves as the primary driver of alpha diversity (Figure 8).

**Figure 8.**
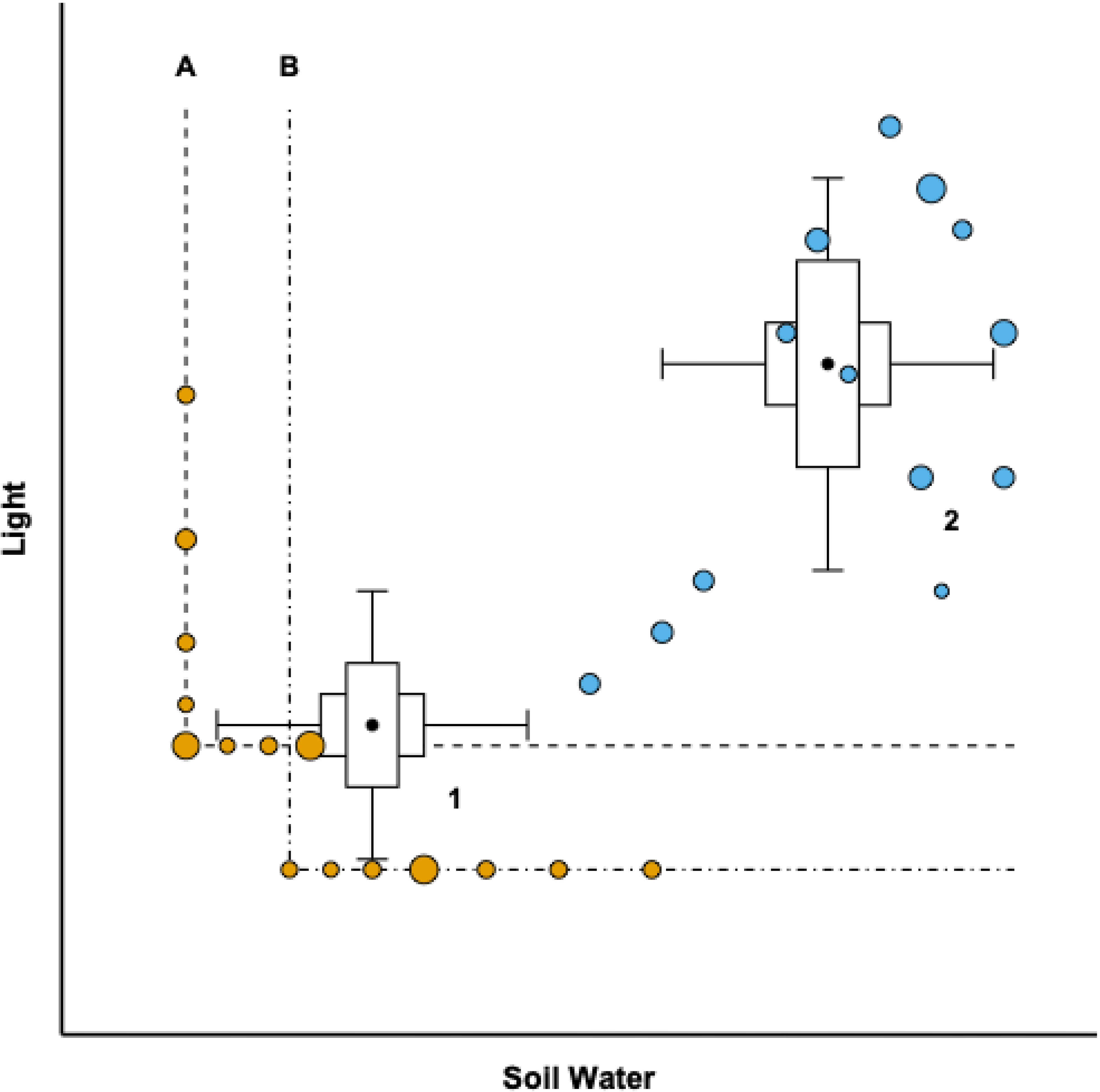
Patchwork Biplot with two species and two metacommunities: 1) stress-governed metacommunity (orange); 2) the same metacommunity governed by disturbance (blue). Here, it was considered that the two resources are consumed or released simultaneously, and the dynamics occur along the diagonal of the mainstream quadrants.

As biomass accumulates in a metacommunity and resource levels (R1 and R2) approach species limiting isoclines, light frequently emerges as the primary limiting factor—particularly in dense tropical forests. In such cases, light availability becomes a critical variable within the Patchwork Biplot framework. Light-limitation stress exerts a powerful structuring force in high-biomass systems. Indeed, Tilman (Tilman, 1988) emphasized light in his biplots, recognizing its central role in R* resource-ratio models. In this example, species B is more shade-tolerant than species A.

### Metacommunities ruled by disturbances transitioning to stress dominance, and vice versa

Consider a watershed affected by a severe hailstorm (Figure 9). At t1, the forest communities are mature and stress-governed, characterized by high biomass and extreme understory resource depletion. Following the storm (t2), these communities transition to a disturbance-governed regime marked by reduced biomass and increased resource availability, particularly light and soil nitrogen. Simulating variations in species richness and patch number under the constraint that plant growth depletes both focal resources reveals distinct temporal trajectories. Comparing metacommunities between t1 (stress-ruled) and t2 (disturbance-governed) shows that trajectories traverse the diagonal Mainstream Quadrants as biomass accumulates or declines.

**Figure 9.**
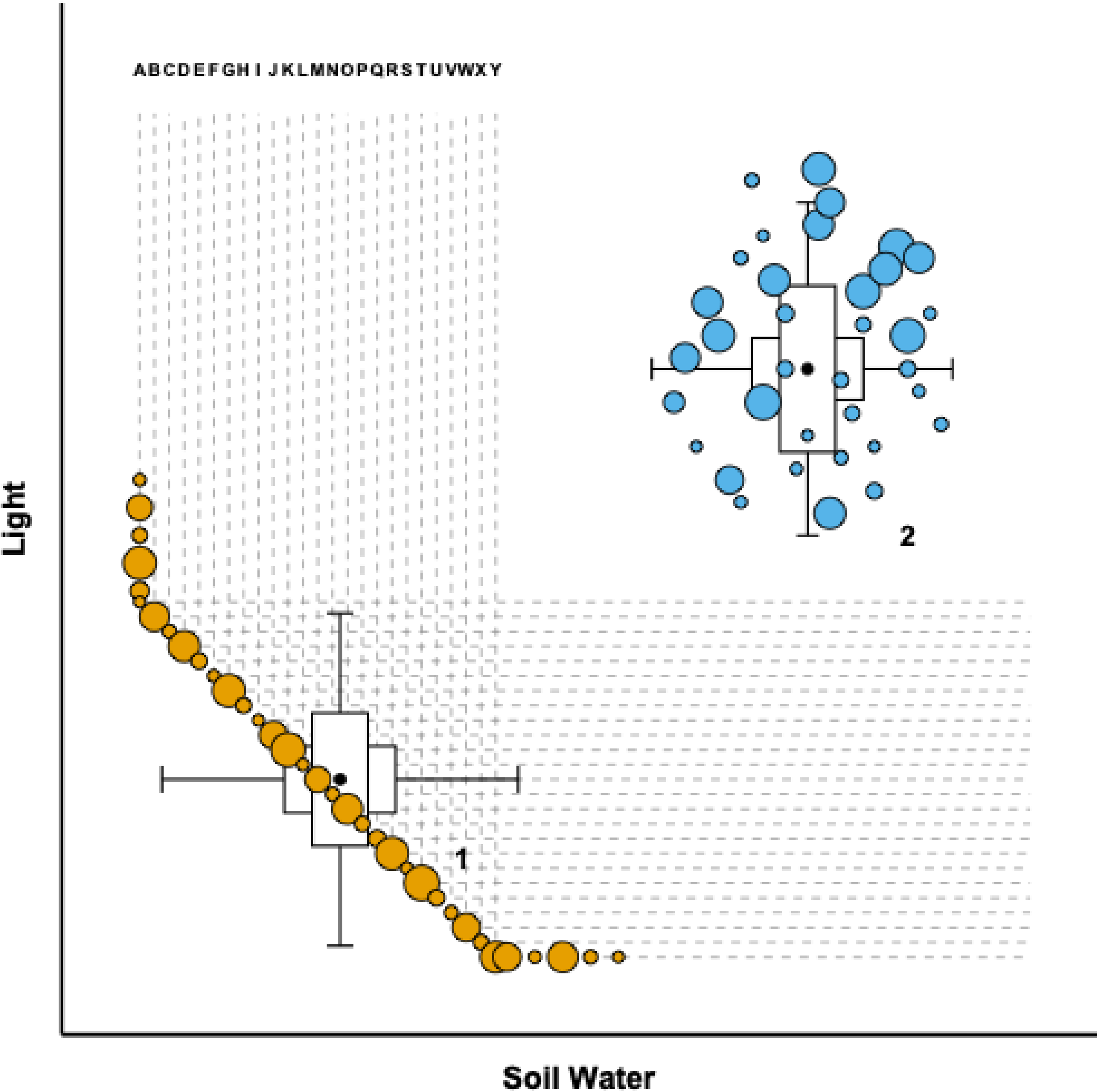
Patchwork Biplot of a metacommunity at two time points (therefore, two metacommunities) with 22 species and 35 communities. 1) Stress-governed metacommunity with communities in orange. 2) Disturbance-governed metacommunity in blue. The concomitant release of water into the soil and light during severe hailstorm occurs without export of these resources, and the resulting dynamics lead the metacommunity to conditions that deviate from the limiting isoclines for the species. Otherwise, interruption or significant decrease in disturbances in 2 can lead the metacommunity towards the isoclines, as exemplified for the orange communities, congruently with what the R* model predicts for communities.

Under high immigration rates, varying species richness yields consistent community compositions across disturbance-governed states (Figure 9). In species- and community-rich scenarios, two or more species can co-occur locally, even if biomass production equilibrium does not converge on a single isocline intersection. Larger community patch sizes (circle areas) increase the likelihood that species from neighboring isoclines will achieve local coexistence. For instance, in a system with 25 species and 35 communities, large-area patches frequently overlap multiple isocline intersections. While this spatial overlap does not guarantee stable multi-species coexistence at the patch level, it enhances the potential for stable coexistence when evaluated at finer spatial scales.

In contrast, anthropogenic disturbances involving net resource export, such as clear-cutting, alter these dynamics. Following clear-cutting, light availability increases dramatically and ceases to be the primary limiting factor as communities transition back toward stress governance. Instead, other resources become limiting, notably soil nitrogen, which is highly mobile and exported alongside organic matter via timber harvesting, biomass burning, and volatilization. Unsurprisingly, Tilman (1988) prioritized nitrogen in his formulation of R* resource-ratio models.

When disturbance involves biomass mortality without resource export, communities shift toward the upper-right quadrant in the biplot. Resources released by decaying biomass move the metacommunity away from resource-limitation-driven species loss, alleviating competitive exclusion (sensu Tilman 1988) and boosting local alpha diversity when high immigration rates are maintained. Conversely, when biomass loss includes resource export, community trajectories traverse the Rare Quadrants: light availability increases, whereas soil nutrients diminish as they are exported alongside lost biomass.

The Patchwork Biplot framework can compare multiple metacommunities or track a single metacommunity over time using intersecting boxplots—for example, comparing a stress-governed state (t1) with a disturbance-governed state (t2) following a widespread disturbance that shifted communities away from isoclines.

Larger community patches hold greater potential for multi-species coexistence, though resolving these micro-dynamics requires spatial downscaling. Because classic R* theory posits that coexistence presupposes biomass equilibrium during interspecific competition, the model can be downscaled to the plant–plant interaction scale where competition actually occurs (Tilman, 1988). If downscaling a hypothetical system of 25 species and 35 communities reveals localized coexistence among four species through pairwise equilibria, the potential for species coexistence in hyperdiverse tropical metacommunities becomes immense, providing a mechanistic basis for how numerous species maintain stable coexistence within a single local community.

In t1, where stress governs communities under high immigration rates, functional trade-offs act as strategy-defining mechanisms that operate alongside simple resource-limitation competition (Figure 9). The Tolerance–Fecundity Trade-off (Muller-Landau, 2010) predicts that small-seeded species generate more offspring, enhancing establishment under favorable conditions, but exhibit lower tolerance to environmental stress compared to large-seeded species. Large-seeded species tolerate high stress, enabling them to persist alongside other stress-tolerant species in high-biomass, resource-depleted communities. Conversely, small-seeded species are more likely to colonize fine-scale resource flushes, such as canopy gaps where nutrients and light temporarily increase. Thus, downscaling reveals that the Tolerance–Fecundity Trade-off can maintain coexistence even when a system reaches a limiting isocline. In this example, circle dimensions correspond to standard sampling plot scales.

Following a large-scale disturbance event, a metacommunity can shift from a stress-governed regime to a disturbance-governed regime, alleviating resource limitations and temporarily relaxing competitive exclusion. Within this disturbance-governed framework, the Competition–Colonization Trade-off explains how species assemble across local communities (Figure 9, blue).

While disturbance generally promotes diversity, intermediate disturbance regimes enhance diversity more effectively than either low or high disturbance levels. This occurs because species specialize along a functional gradient ranging from low-disturbance specialists (strong competitors) to high-disturbance specialists (efficient colonizers). Intermediate disturbances enable the coexistence of species from both ends of this spectrum, thereby maximizing local alpha diversity. Downscaling to intra-community scales demonstrates how fine-scale spatial variation in disturbance further supports coexistence. In the same way, downscaling to intracommunity scale the fine-scale variation in stress further supports coexistence (Figure 10). Larger community patches—represented by larger circle dimensions—encompass greater internal variation, thereby expanding the potential for localized multi-species coexistence.

**Figure 10.**
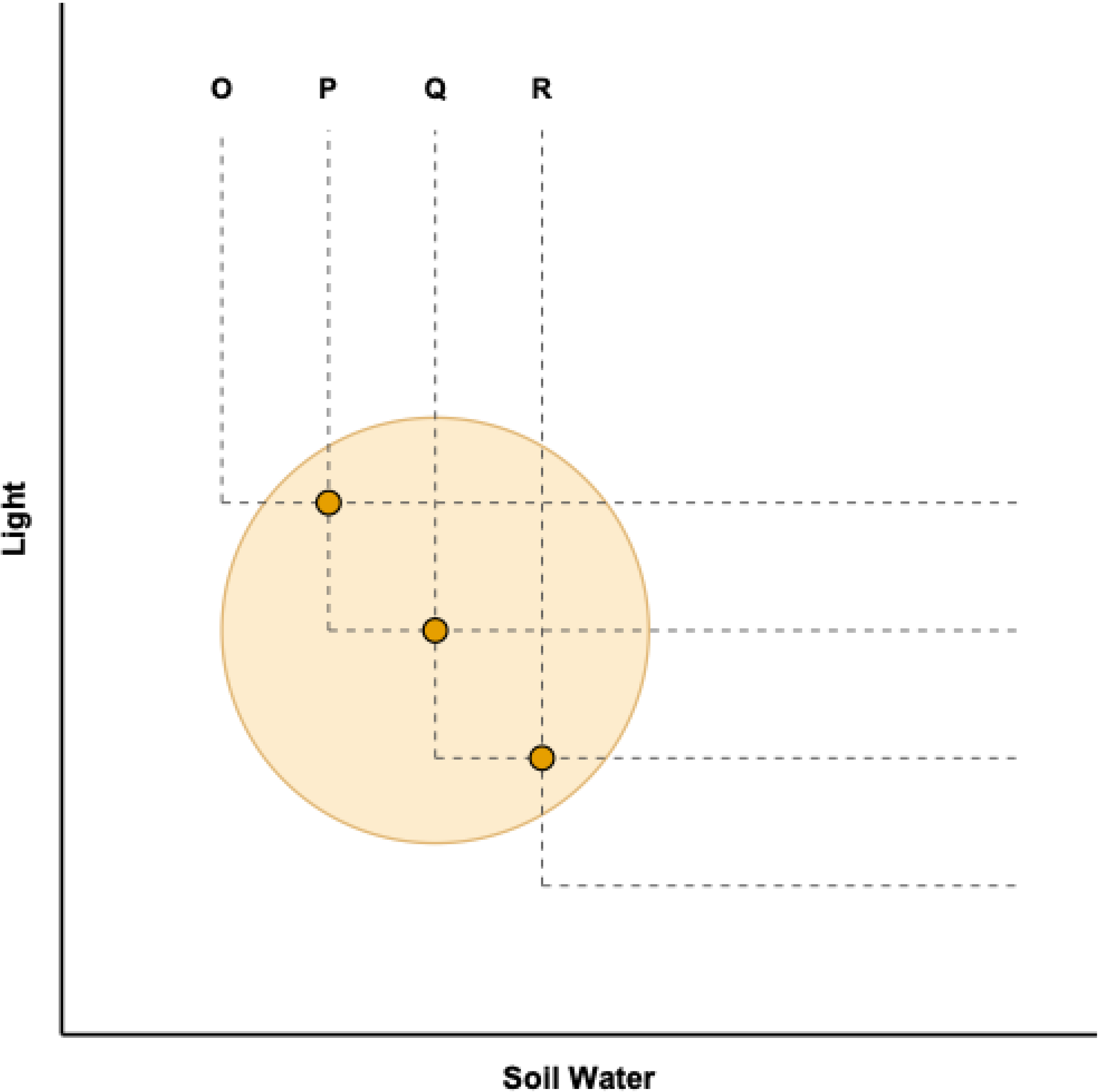
A detail of the Patchwork Biplot. A community (large orange circle) of the Figure 11 when analyzed at a smaller scale can register the effects of different granulation due to soil drainage or microtopography. In this case, even with species competing directly for light and water in the soil, coexistence can be achieved by several species through the granulation of the smaller scale. The larger the area of the community, the greater the potential for variation in resources at smaller scales.

## Materials and Methods - Case Study

### Paraopeba Reserve and vegetation sampling

The 203-hectare Paraopeba Reserve features five soil types, with 150 hectares consisting of native Cerrado vegetation situated within the core distribution of the biome. Although the Cerrado is a fire-prone ecosystem, similar to Mediterranean maquis and scrublands (Meira-Neto et al., 2011), the reserve has been strictly protected from fire since 1963. Located in the state of Minas Gerais (19°16′19″ S, 44°24′06″ W), its elevation ranges from 734 m above sea level (ASL) in the south to 761 m ASL in the north. Vegetation sampling data were drawn from plots in each of the soil types of Paraopeba Reserve (Tolentino, 2011). In each of the five soil types, five 20 x 20 m plots were settled, totaling 1 hectare, in which woody plants with stems of 10 cm or more at ground level were sampled.

### Soils and total nitrogen

Twenty-five 20 × 20 m square plots (400 m² each) were established along transects, yielding a cumulative sampled area of 1 hectare. The sampling design stratified the reserve across five distinct soil environments, allocating five plots to each soil type: Yellow Cambisol, Yellow Latosol, Red-Yellow Latosol, Dystrophic Red Latosol, and Mesotrophic Red Latosol (Neri et al., 2013).

Soil was sampled from the 0–20 cm depth layer using a composite sampling approach, with subsamples combined and thoroughly homogenized for each plot. In the laboratory, samples were air-dried, disaggregated, and passed through a 2 mm mesh sieve to obtain the Air-Dried Fine Earth fraction.

To determine total nitrogen content, the ADFE fraction was analyzed via sulfuric acid digestion (Micro-Kjeldahl method) according to standard Embrapa protocols (Teixeira et al., 2017). Soil samples were heated in a digestion block with concentrated sulfuric acid and a catalytic mixture to oxidize organic matter and convert all organic nitrogen into ammonium ions. After cooling, the digestate was alkalized with a concentrated sodium hydroxide solution and steam-distilled to release gaseous ammonia. The liberated ammonia was captured in a boric acid receiving solution containing a mixed indicator and quantified by titration against a standardized hydrochloric or sulfuric acid solution, yielding total soil nitrogen content expressed in dag kg⁻¹.

### Hemispherical photographs and canopy openness

Canopy openness was sampled at the center of each plot during the rainy season (January–February 2010). Hemispherical photographs were captured under diffuse light conditions (overcast skies) using a Nikon Coolpix 5700 digital camera fitted with a Nikon UR-E12 extender and a Nikon FC-E9 fisheye lens. The camera was mounted on a tripod 1.5 m above the ground, oriented toward the zenith, with the top of the frame aligned with magnetic north.

Images were analyzed using Gap Light Analyzer software (GLA, version 2.0) (Frazer et al., 1999). The software generated a binary (black-and-white) mask for each photograph to distinguish canopy cover from gaps within the hemispherical projection. It automatically calculated percent canopy openness—defined as the proportion of unobstructed sky relative to the total area of the hemispherical projection, representing the inverse of canopy cover.

## Results and Discussion - Case Study

The soil nitrogen and light availability is the most explored example of resource-ratio model in Tilman (1988). Because the Paraopeba Reserve has been effectively protected from severe disturbance for several decades, these communities would be stress-ruled. Furthermore, given the absence of dispersal limitation within the reserve, community vectors are expected to remain parallel.

Cerrado dynamics under contrasting fire regimes were evaluated by Belmok et al. (Belmok et al., 2019) across recently burned and unburned plots in the IBGE Ecological Reserve in Brasília. In recently burned Cerrado, mean total nitrogen (N) was below 0.07 dag kg⁻¹ (ranging from 0.05 to 0.08 dag kg⁻¹). Unburned plots exhibited mean N values of 0.13 dag kg⁻¹ (ranging from 0.11 to 0.15 dag kg⁻¹), both lower than the minimum value recorded in our protected reserve (0.157 dag kg⁻¹). Fire volatilizes most aboveground biomass nitrogen, leaving only a fraction of total N stocks in post-fire ash (Pivello and Coutinho 1992). Consequently, despite a temporary surge in surface NH₄⁺ availability following a burn, annual N cycling in the Cerrado drops sharply post-fire (Nardoto & Bustamante, 2003). Accordingly, minimum soil N levels at the Paraopeba Reserve exceed those reported for Cerrado sites with recent fire histories, a widespread baseline given the fire-prone nature of Cerrado ecosystems (Ribeiro & Walter, 1998; Hoffmann, 2002; Simon et al., 2009). In the Patchwork Biplot for the Paraopeba Reserve, canopy openness values indicate strongly light-limited conditions, ranging from 7.5% to 52.13%. This range shows minimal overlap with the 50–80% canopy openness (corresponding to 20-50% canopy cover) typical of Cerrado vegetation broadly (Ribeiro & Walter, 1998).

In the Patchwork Biplot for the Paraopeba Reserve of this case study, we incorporated assumed data (represented by the blue ellipse) using soil nitrogen (N) ranges from the IBGE Reserve Cerrado under recently burned and unburned conditions (Belmok et al., 2019), alongside typical canopy openness ranges for the Cerrado biome (Ribeiro & Walter, 1998). This comparison indicates that the Paraopeba Reserve plots experience light limitation (stress), but not N limitation. Consequently, all observed, non-simulated plots fall outside the blue ellipse and are designated in orange.

As the Patchwork Biplot isoclines do not necessarily denote zero-net-growth boundaries or absolute niche limits for these species, in this case study it represents the most limiting situation of a species relatively to a resource in a metacommunity. The blue ellipse is beyond the nitrogen isoclines and represents the historical baseline of the plots prior to active fire protection in the reserve (Figure 11). Therefore, stress-ruled communities (orange) align along horizontal isoclines in the biplot, denoting limiting isoclines. In contrast, communities recovering from past disturbances (blue) exhibit net trajectories toward these horizontal canopy openness isoclines as canopy openness decreases, shading increases and soil N accumulates (Figures 12, S1 and Table S1 and S2). The prolonged absence of disturbance reduces canopy openness and allows soil N stocks to build up, mirroring dynamics reported for unburned Cerrado sites (Nardoto & Bustamante, 2003; Belmok et al., 2019).

**Figure 11.**
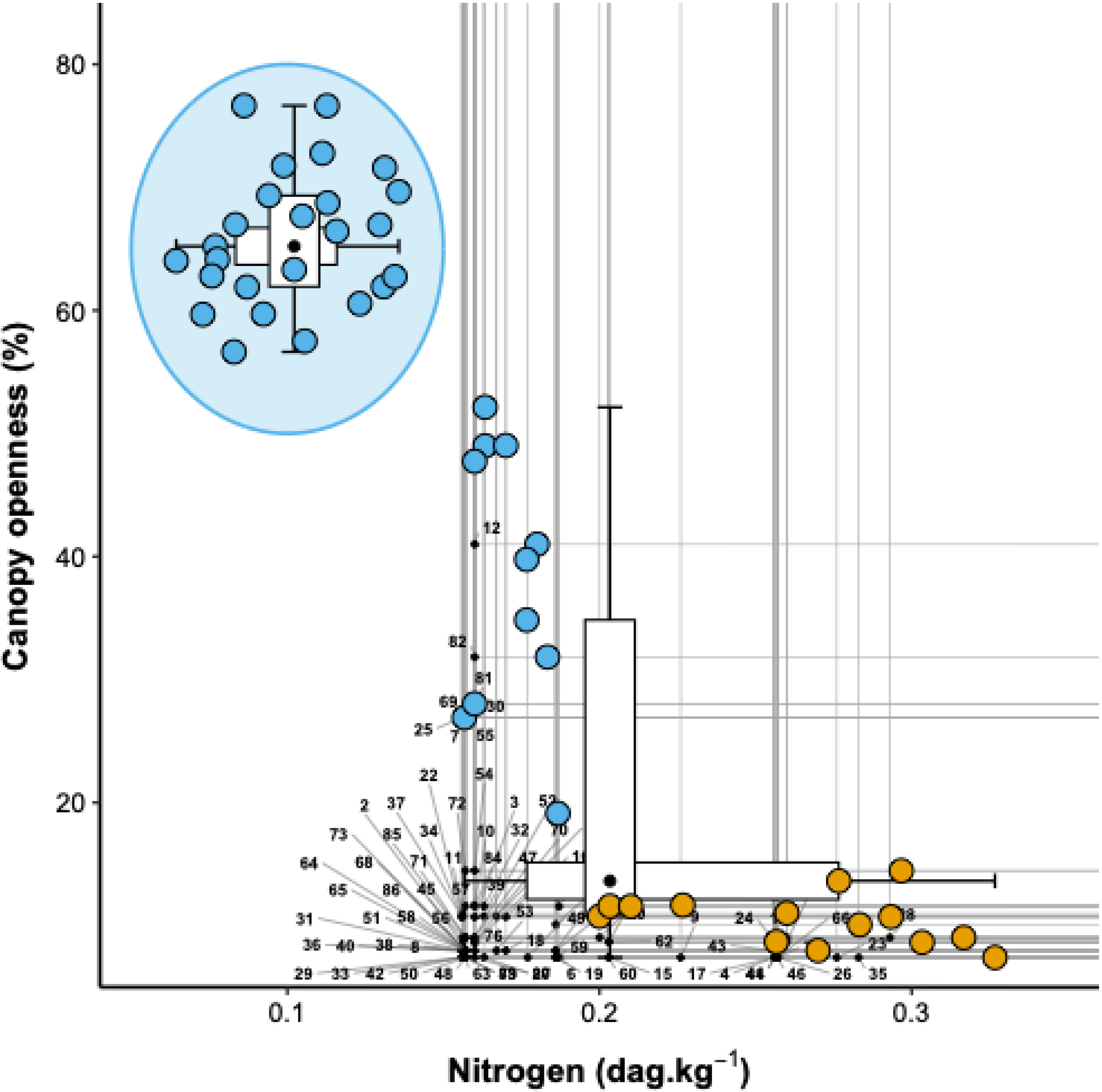
Patchwork Biplot of plots in the Paraopeba Reserve Cerrado. Inside the blue ellipse: Simulation of the communities of the Paroapeba Reserve when assuming N values reported by Belmok et al. (2019) for Cerrados with a fire regime in recent decades, and canopy openness values by (Ribeiro & Walter, 1998) for the expected range of variation for the Cerrado in general. Outside the blue ellipse: in the upper left quadrant of the Patchwork Biplot, the communities may still retain some past effect of disturbances prior to the protection of the Paroapeba Reserve, therefore in this figure they are in blue, indicating that they are not yet predominantly governed by stress; In the lower right quadrant, communities are organized along canopy openness contours, suggesting sliding along the contours as predicted in Figure 7. Past disturbances in regimes similar to those of the IBGE Reserve (large blue circle) increase canopy openness, increasing illumination at ground level and decreasing shading stresses. Communities in orange are under light limitation and governed by stress, in the lower rare quadrant. As the canopy openness values of N were obtained only for the Paraopeba Reserve; therefore, the communities in the blue ellipse did not define the isoclines.

**Figure 12.**
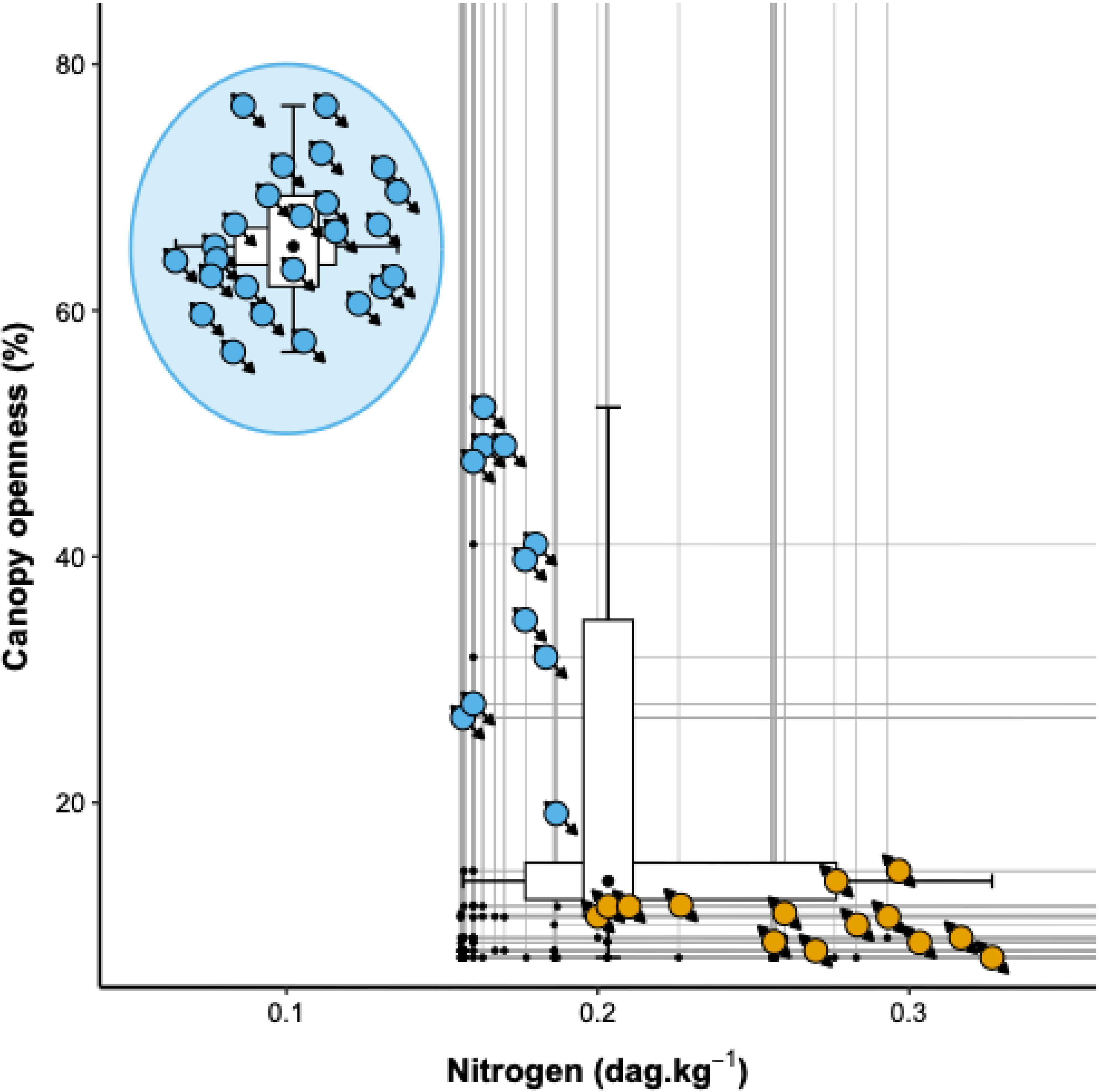
The Cerrado metacommunity of the Paraopeba Reserve appears here at two different times, with dynamics diagonally across the rare quadrants. The metacommunity at the top and inside the blue ellipse is a simulation of the Paraopeba Reserve metacommunity before fire disturbance protection. Outside the blue ellipse, the blue communities with net vectors pointing towards the horizontal canopy openness isoclines are communities that still have light that can be extinguished, possibly due to increased soil N levels. In orange, the horizontal arrangement of the communities along the canopy openness isoclines suggests that oscillations in canopy openness values and the continuous increase in N slide the communities to the right. Note the different vector sizes in the different situations (see Figure 7). As the canopy openness, the N values were obtained only for the Paraopeba Reserve, the communities in the blue ellipse did not define the isoclines.

As soil N increases, community trajectories traverse the Rare Quadrants, with disturbed plots (blue) passing through the upper Rare Quadrant and protected plots (orange) occupying the lower Rare Quadrant (but see also Figures S2 and S3, for an example with Mainstream Quadrants directions). These trajectories eventually settle along species isoclines at minimum light tolerance thresholds, potentially oscillating rightward along these horizontal boundaries (Figure 7). Notably, species with higher canopy openness isoclines face exclusion not from a single superior competitor, but from any species exhibiting greater shade tolerance. Consequently, light-driven competitive exclusion nowadays can be a pervasive structuring force across the Paraopeba Reserve.

In the Paraopeba Cerrado comparatively to the position of the metacommunity before the fire supression, metacommunity dynamics suggest that intensification of fire regimes can induce nitrogen (N) stress. The Patchwork Biplot (Figure 12) indicates that competitive exclusion intensifies under both complete disturbance suppression, with light as limiting factor, and extreme disturbance regimes, with N as limiting factor.

It is not clear if the species that are depleting nitrogen and canopy openness concurrently are able to deplete the resources up to their vertices. In some cases resources interact in a way that high availability of one resource enables high tolerance of another resource limitation (Tilman, 1988). That would be the same case of leaves with high investment of N in photosynthetic enzymes and pigments that improve light harvesting per mass unit of tissues (Reich et al., 1992, 1999).

Our results demonstrate how the Patchwork Biplot serves as an objective tool, for instance, to prioritize species vulnerable to competitive exclusion across local communities and the broader metacommunity. Replacing subjective classification schemes, this model-based graphical framework clarifies species-specific vulnerabilities while identifying targeted management actions to prevent local extirpations. In our case study, declining canopy openness highlights the heightened vulnerability of *Byrsonima verbascifolia*, *Tocoyena formosa*, *Tibouchina granulosa*, *Erythroxylum deciduum*, *Palicourea rigida*, *Didymopanax macrocarpus*, *Salvertia convallariodora*, and *Baccharis platypoda*, establishing a prioritized hierarchy for conservation intervention (Figure 11, Table 1). To prevent competitive exclusion across this metacommunity, management should prioritize increasing canopy openness by reducing dense woody vegetation cover. Specific intervention strategies, such as prescribed burning or alternative management practices, should be formally evaluated and incorporated into the reserve’s management plan.

**Table 1.**
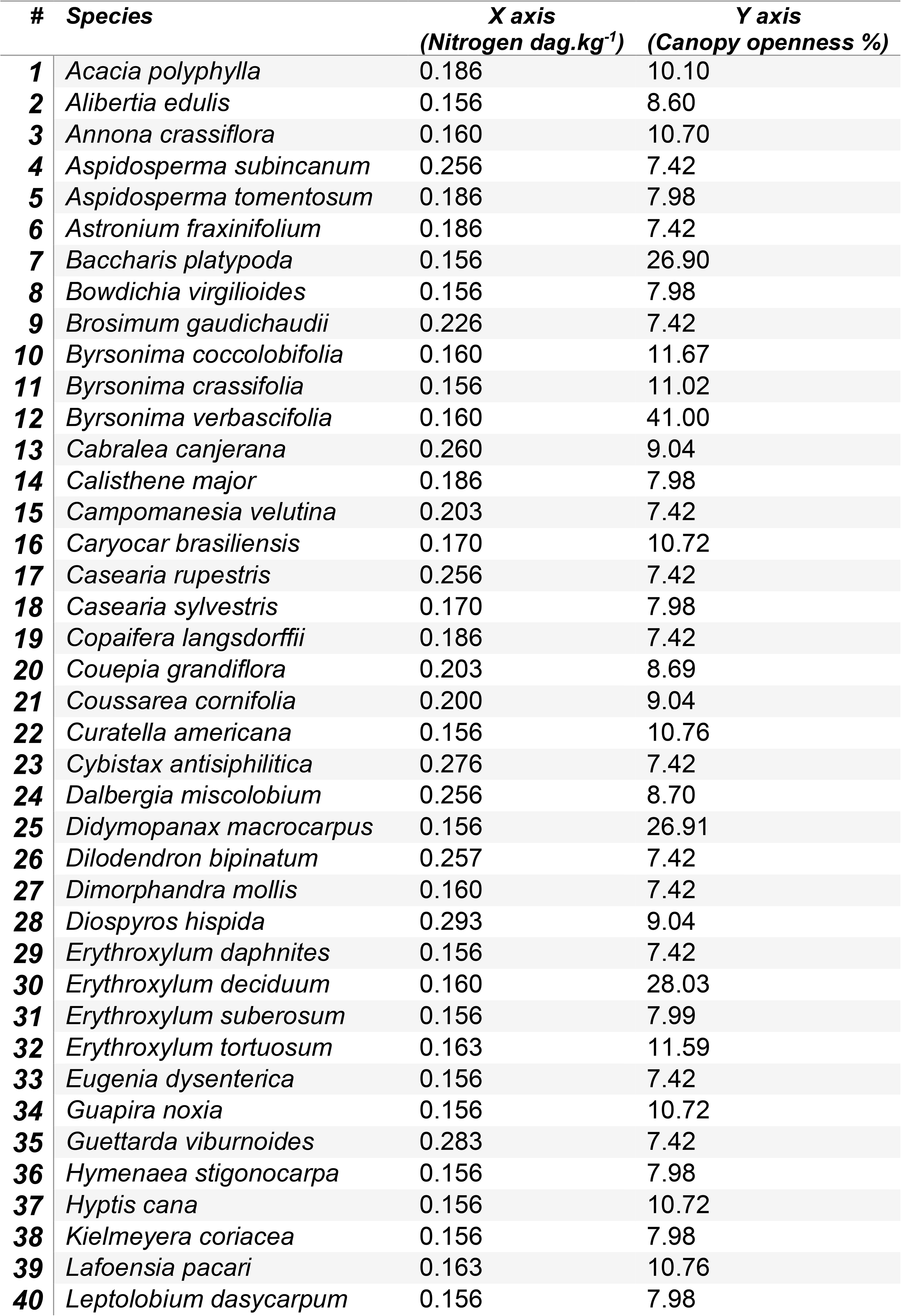
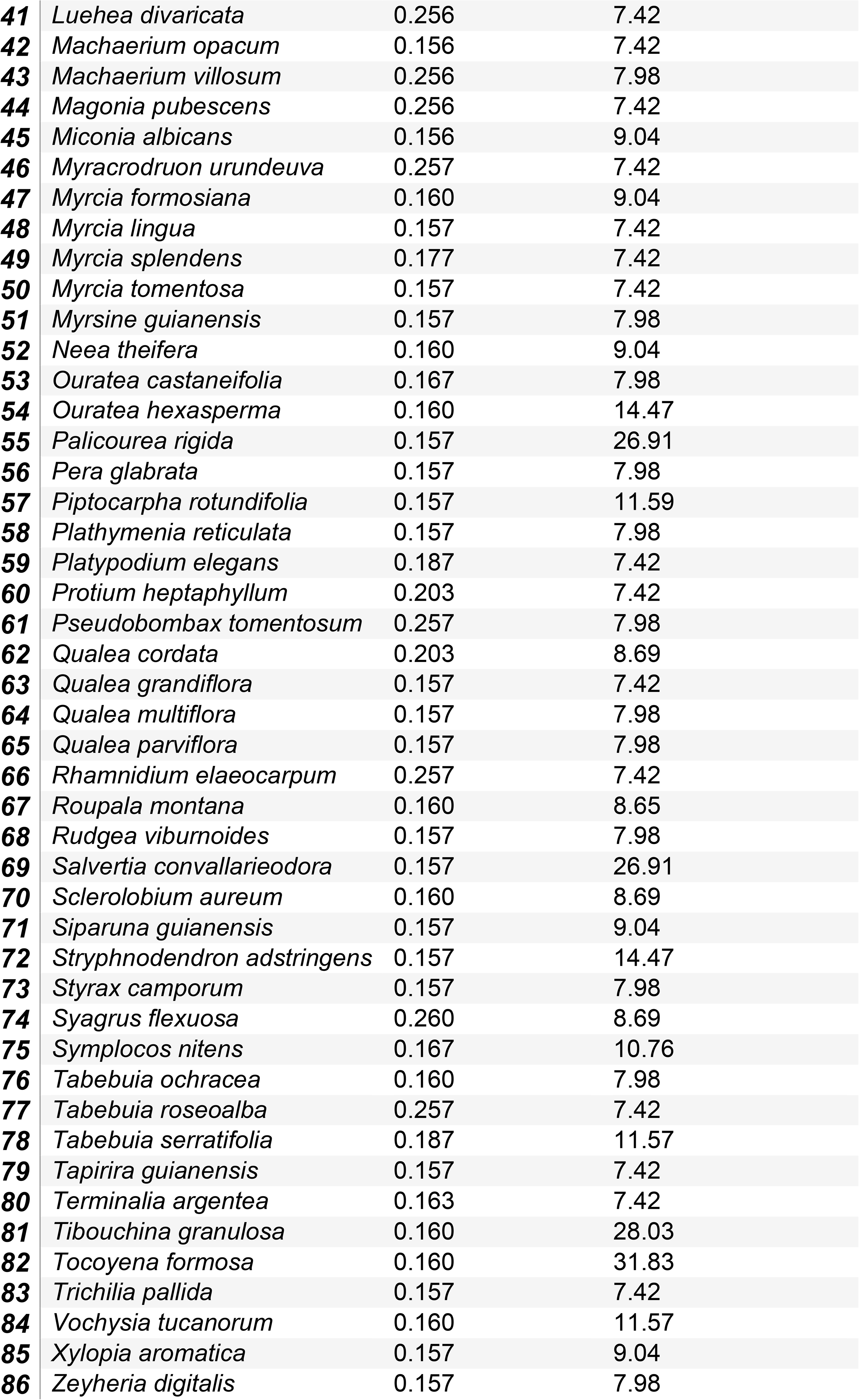
Species references and x, y coordinates of species in the Patchwork Biplot of the case study. Only species with sampled individuals in at least three communities were considered for isoclines.

| # | Species | X axis<br>(Nitrogen dag.kg<sup>-1</sup>) | Y axis<br>(Canopy openness %) |
| --- | --- | --- | --- |
| 1 | *Acacia polyphylla* | 0.186 | 10.10 |
| 2 | *Alibertia edulis* | 0.156 | 8.60 |
| 3 | *Annona crassiflora* | 0.160 | 10.70 |
| 4 | *Aspidosperma subincanum* | 0.256 | 7.42 |
| 5 | *Aspidosperma tomentosum* | 0.186 | 7.98 |
| 6 | *Astronium fraxinifolium* | 0.186 | 7.42 |
| 7 | *Baccharis platypoda* | 0.156 | 26.90 |
| 8 | *Bowdichia virgilioides* | 0.156 | 7.98 |
| 9 | *Brosimum gaudichaudii* | 0.226 | 7.42 |
| 10 | *Byrsonima coccolobifolia* | 0.160 | 11.67 |
| 11 | *Byrsonima crassifolia* | 0.156 | 11.02 |
| 12 | *Byrsonima verbascifolia* | 0.160 | 41.00 |
| 13 | *Cabralea canjerana* | 0.260 | 9.04 |
| 14 | *Calisthene major* | 0.186 | 7.98 |
| 15 | *Campomanesia velutina* | 0.203 | 7.42 |
| 16 | *Caryocar brasiliensis* | 0.170 | 10.72 |
| 17 | *Casearia rupestris* | 0.256 | 7.42 |
| 18 | *Casearia sylvestris* | 0.170 | 7.98 |
| 19 | *Copaifera langsdorffii* | 0.186 | 7.42 |
| 20 | *Couepia grandiflora* | 0.203 | 8.69 |
| 21 | *Coussarea cornifolia* | 0.200 | 9.04 |
| 22 | *Curatella americana* | 0.156 | 10.76 |
| 23 | *Cybistax antisiphilitica* | 0.276 | 7.42 |
| 24 | *Dalbergia miscolobium* | 0.256 | 8.70 |
| 25 | *Didymopanax macrocarpus* | 0.156 | 26.91 |
| 26 | *Dilodendron bipinatum* | 0.257 | 7.42 |
| 27 | *Dimorphandra mollis* | 0.160 | 7.42 |
| 28 | *Diospyros hispida* | 0.293 | 9.04 |
| 29 | *Erythroxylum daphnites* | 0.156 | 7.42 |
| 30 | *Erythroxylum deciduum* | 0.160 | 28.03 |
| 31 | *Erythroxylum suberosum* | 0.156 | 7.99 |
| 32 | *Erythroxylum tortuosum* | 0.163 | 11.59 |
| 33 | *Eugenia dysenterica* | 0.156 | 7.42 |
| 34 | *Guapira noxia* | 0.156 | 10.72 |
| 35 | *Guettarda viburnoides* | 0.283 | 7.42 |
| 36 | *Hymenaea stigonocarpa* | 0.156 | 7.98 |
| 37 | *Hyptis cana* | 0.156 | 10.72 |
| 38 | *Kielmeyera coriacea* | 0.156 | 7.98 |
| 39 | *Lafoensia pacari* | 0.163 | 10.76 |
| 40 | *Leptolobium dasycarpum* | 0.156 | 7.98 |
| 41 | *Luehea divaricata* | 0.256 | 7.42 |
| 42 | *Machaerium opacum* | 0.156 | 7.42 |
| 43 | *Machaerium villosum* | 0.256 | 7.98 |
| 44 | *Magonia pubescens* | 0.256 | 7.42 |
| 45 | *Miconia albicans* | 0.156 | 9.04 |
| 46 | *Myracrodruon urundeuva* | 0.257 | 7.42 |
| 47 | *Myrcia formosiana* | 0.160 | 9.04 |
| 48 | *Myrcia lingua* | 0.157 | 7.42 |
| 49 | *Myrcia splendens* | 0.177 | 7.42 |
| 50 | *Myrcia tomentosa* | 0.157 | 7.42 |
| 51 | *Myrsine guianensis* | 0.157 | 7.98 |
| 52 | *Neea theifera* | 0.160 | 9.04 |
| 53 | *Ouratea castaneifolia* | 0.167 | 7.98 |
| 54 | *Ouratea hexasperma* | 0.160 | 14.47 |
| 55 | *Palicourea rigida* | 0.157 | 26.91 |
| 56 | *Pera glabrata* | 0.157 | 7.98 |
| 57 | *Piptocarpha rotundifolia* | 0.157 | 11.59 |
| 58 | *Plathymenia reticulata* | 0.157 | 7.98 |
| 59 | *Platypodium elegans* | 0.187 | 7.42 |
| 60 | *Protium heptaphyllum* | 0.203 | 7.42 |
| 61 | *Pseudobombax tomentosum* | 0.257 | 7.98 |
| 62 | *Qualea cordata* | 0.203 | 8.69 |
| 63 | *Qualea grandiflora* | 0.157 | 7.42 |
| 64 | *Qualea multiflora* | 0.157 | 7.98 |
| 65 | *Qualea parviflora* | 0.157 | 7.98 |
| 66 | *Rhamnidium elaeocarpum* | 0.257 | 7.42 |
| 67 | *Roupala montana* | 0.160 | 8.65 |
| 68 | *Rudgea viburnoides* | 0.157 | 7.98 |
| 69 | *Salvertia convallarieodora* | 0.157 | 26.91 |
| 70 | *Sclerolobium aureum* | 0.160 | 8.69 |
| 71 | *Siparuna guianensis* | 0.157 | 9.04 |
| 72 | *Stryphnodendron adstringens* | 0.157 | 14.47 |
| 73 | *Styrax camporum* | 0.157 | 7.98 |
| 74 | *Syagrus flexuosa* | 0.260 | 8.69 |
| 75 | *Symplocos nitens* | 0.167 | 10.76 |
| 76 | *Tabebuia ochracea* | 0.160 | 7.98 |
| 77 | *Tabebuia roseoalba* | 0.257 | 7.42 |
| 78 | *Tabebuia serratifolia* | 0.187 | 11.57 |
| 79 | *Tapirira guianensis* | 0.157 | 7.42 |
| 80 | *Terminalia argentea* | 0.163 | 7.42 |
| 81 | *Tibouchina granulosa* | 0.160 | 28.03 |
| 82 | *Tocoyena formosa* | 0.160 | 31.83 |
| 83 | *Trichilia pallida* | 0.157 | 7.42 |
| 84 | *Vochysia tucanorum* | 0.160 | 11.57 |
| 85 | *Xylopia aromatica* | 0.157 | 9.04 |
| 86 | *Zeyheria digitalis* | 0.157 | 7.98 |

Other example is that, in the case of fire regime implementation as a management tool, species with isoclines suggesting higher nitrogen requirements are possibly experiencing colonization across the Paraopeba Reserve following disturbance suppression. Consequently, these species would be disproportionately vulnerable to intensified disturbance and soil nitrogen export. In order of priority, these include *Cybistax antisyphilitica*, *Dilodendron bipinnatum*, *Diospyros hispida*, *Guettarda viburnoides*, *Aspidosperma subincanum*, *Casearia rupestris*, *Dalbergia miscolobium*, *Machaerium villosum*, *Magonia pubescens*, *Myracrodruon urundeuva*, and *Protium heptaphyllum* (Figure 11, Table 1).

Another example of the MetcommR and the Patchwork biplot as tools is that the Paraopeba Reserve is in a Global Change context in which we could predict changes in stresses, disturbances and connectivity. Considering stresses, if the main resulting stress is an increased drought intensity or duration, increased nitrogen contents and vegetation cover would mitigate the water stress (Siemann & Rogers, 2003; Meira-Neto, et al., 2018; Meira-Neto, et al., 2018). Considering disturbances, if the fire disturbance increases, the nitrogen possibly becomes the main limiting resource surpassing the canopy openness as stronger stressing factor (Nardoto & Bustamante, 2003; Belmok et al., 2019). Considering connectivity, if the isolation of the Paraopeba Reserve increases in the landscape, the less predictable is the species composition and the less predictable is the nitrogen x canopy openness dynamics in the Patchwork Biplot (Hubbell, 2001; Leibold et al., 2004, 2017).

## Acknowledgements

The author thank the Conselho Nacional de Desenvolvimento Científico e Tecnológico (CNPq) - fellowship 304386/2024-3, FAPEMIG-Botany Graduate Program grant, CAPES - Botany Graduate Program PROAP. The author also thank Glaucia Soares Tolentino, Maria Carolina Alves Nunes da Silva, Marcia Nascimento assistance with fieldwork and laboratory work, FLONA de Paraopeba staff for supporting fieldwork.

---

**Figure S1.**
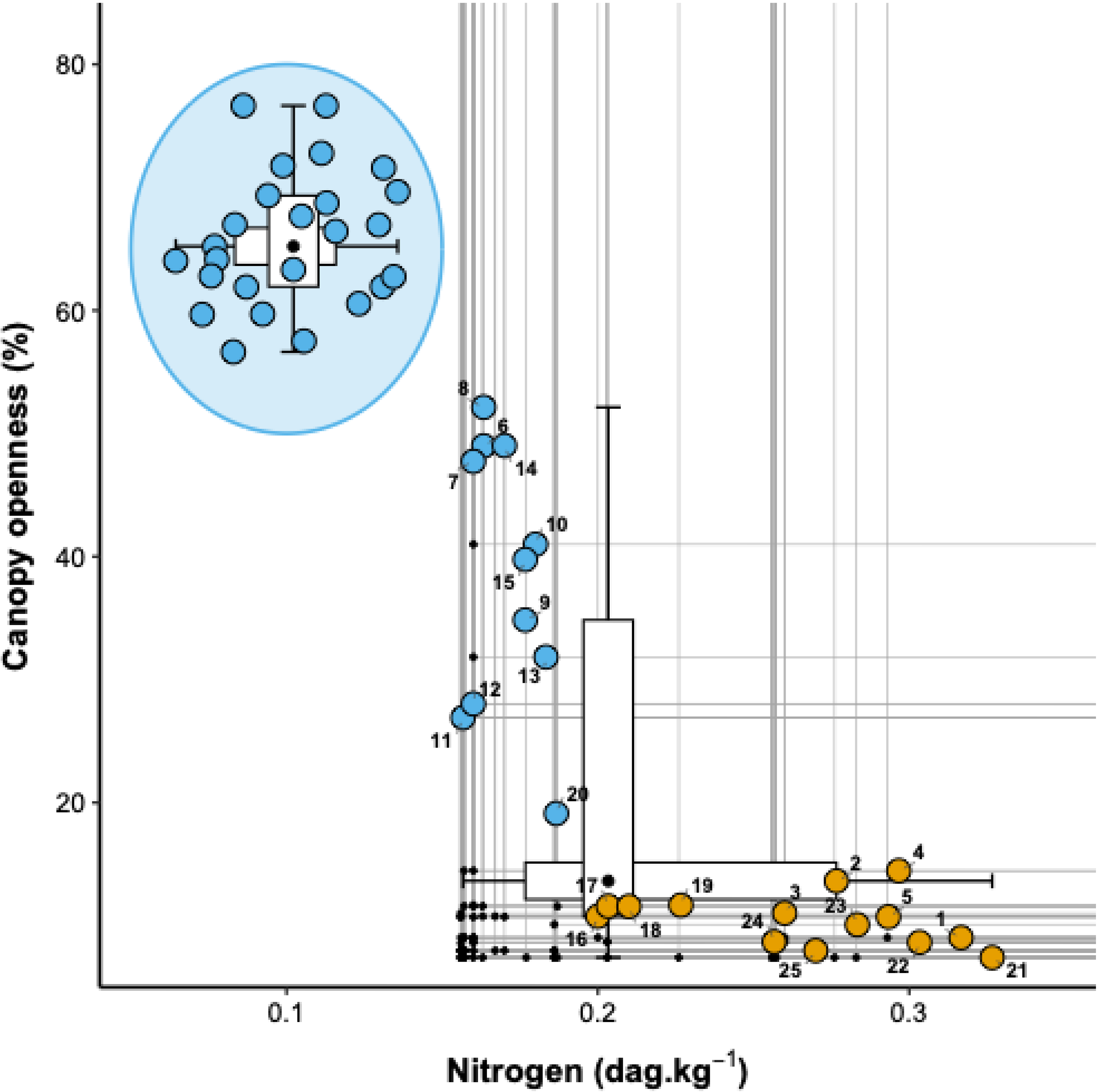
Patchwork Biplot of plots in the Paraopeba Reserve Cerrado. Inside the blue ellipse: Simulation of the communities of the Paroapeba Reserve assuming N values reported by Belmok et al. (2019) for Cerrados with a fire regime in recent decades, and canopy openness values by (Ribeiro & Walter, 1998) for the expected range of variation for the Cerrado in general. Outside the blue ellipse: in the upper left quadrant of the Patchwork Biplot, the communities may still retain some past effect of disturbances prior to the protection of the Paroapeba Reserve, therefore in this figure they are in blue, indicating that they are not yet predominantly governed by stress; In the lower right quadrant, communities are organized along canopy openness contours, suggesting sliding along the contours as predicted in Figure 8. Past disturbances in regimes similar to those of the IBGE Reserve (large blue circle) increase canopy openness, increasing illumination at ground level and decreasing shading stresses. Communities in orange are under light limitation and governed by stress, in the lower rare quadrant. As the canopy openness, the values of N were obtained only for the Paraopeba Reserve; therefore, the communities in the blue ellipse did not define the isoclines.

**Figure S2.**
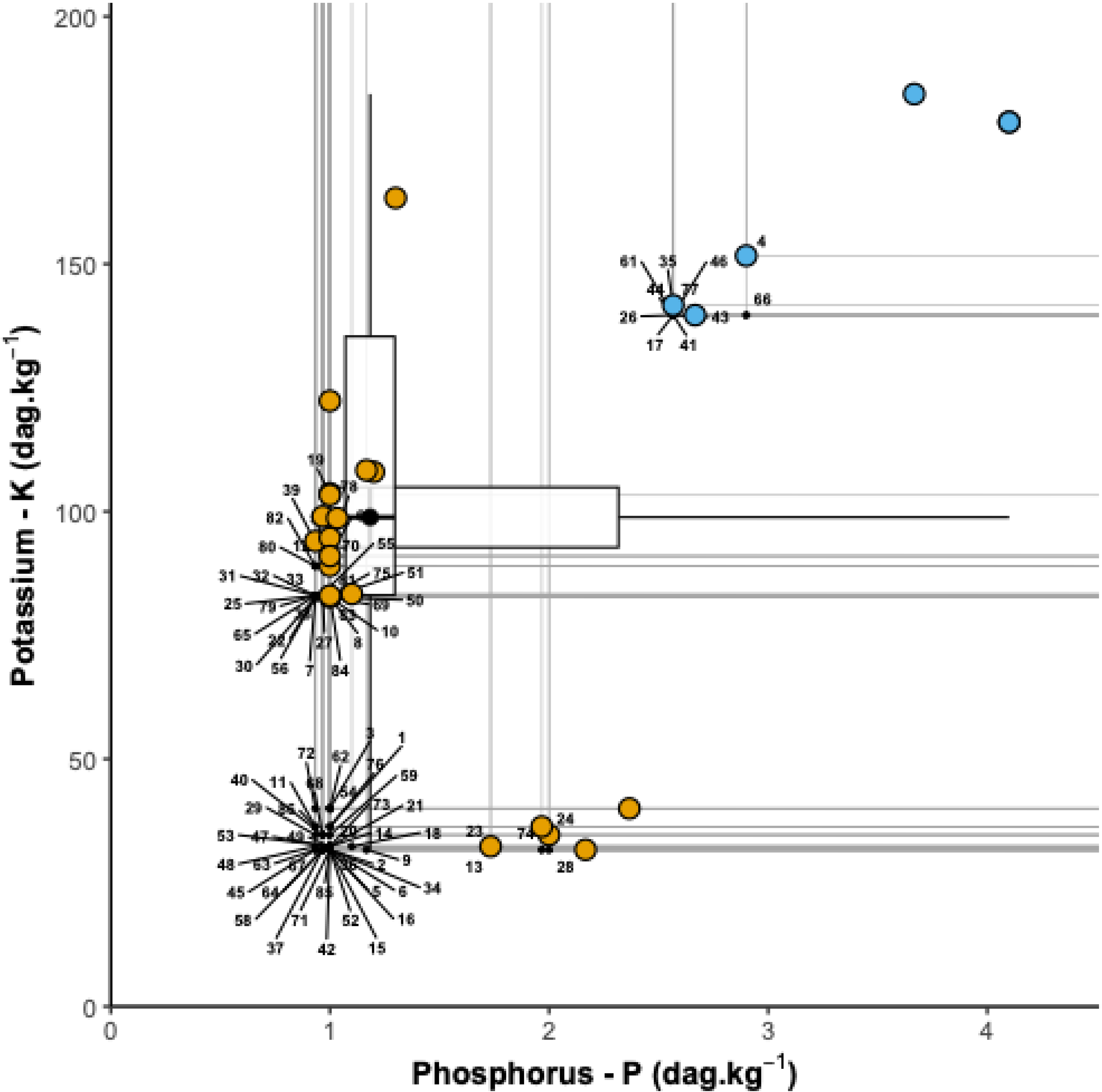
Patchwork Biplot of plots in the Paraopeba Reserve Cerrado with numbers indicating species of the Table S1. In the upper right quadrant (Mainstream Quadrant) of the Patchwork Biplot, the communities may still retain some past effect of disturbances prior to the protection of the Paroapeba Reserve, therefore in this figure they are in blue, indicating that they are not yet predominantly governed by stress; In the lower right quadrant and in the left, communities are organized along phosphorus and potassium contours. Communities in orange are under phosphorus or potassium limitation and governed by stress.

**Figure S3.**
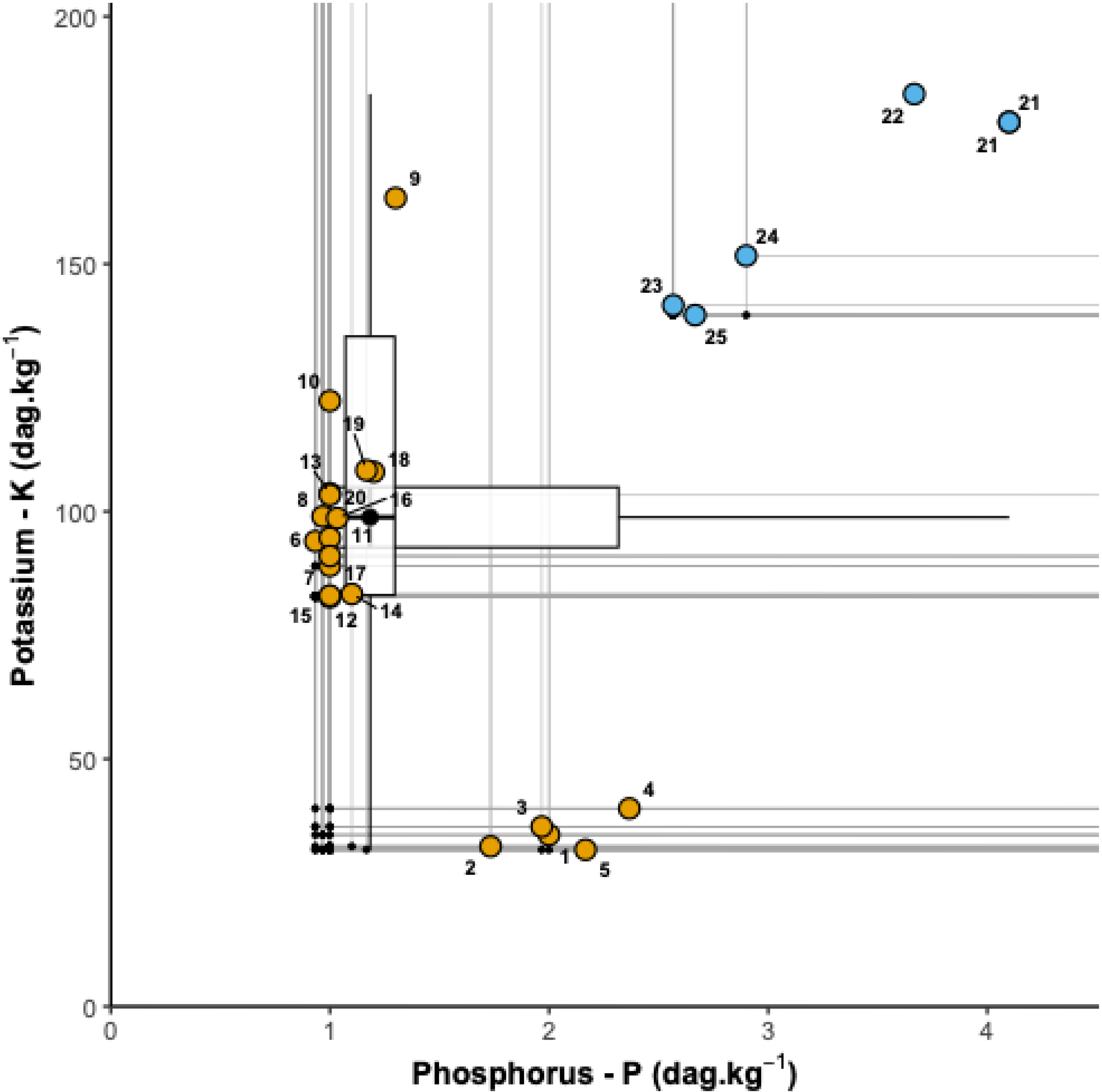
Patchwork Biplot of plots in the Paraopeba Reserve Cerrado with numbers indicating the plots. In the upper right quadrant (Mainstream Quadrant) of the Patchwork Biplot, the communities may still retain some past effect of disturbances prior to the protection of the Paroapeba Reserve, therefore in this figure they are in blue, indicating that they are not yet predominantly governed by stress; In the lower right quadrant and in the left, communities are organized along phosphorus and potassium contours. Communities in orange are under phosphorus or potassium limitation and governed by stress.

**Table S1.**
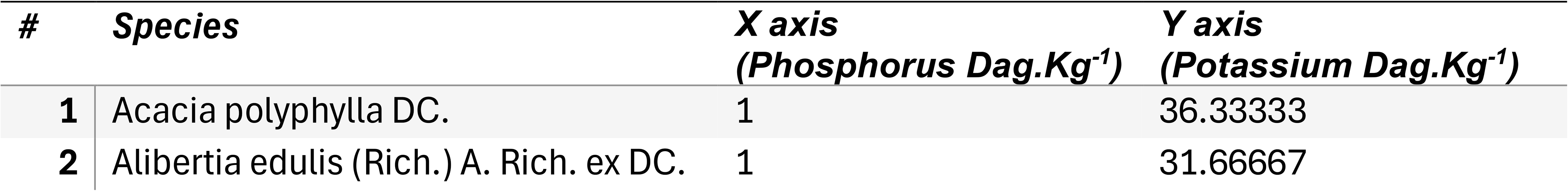
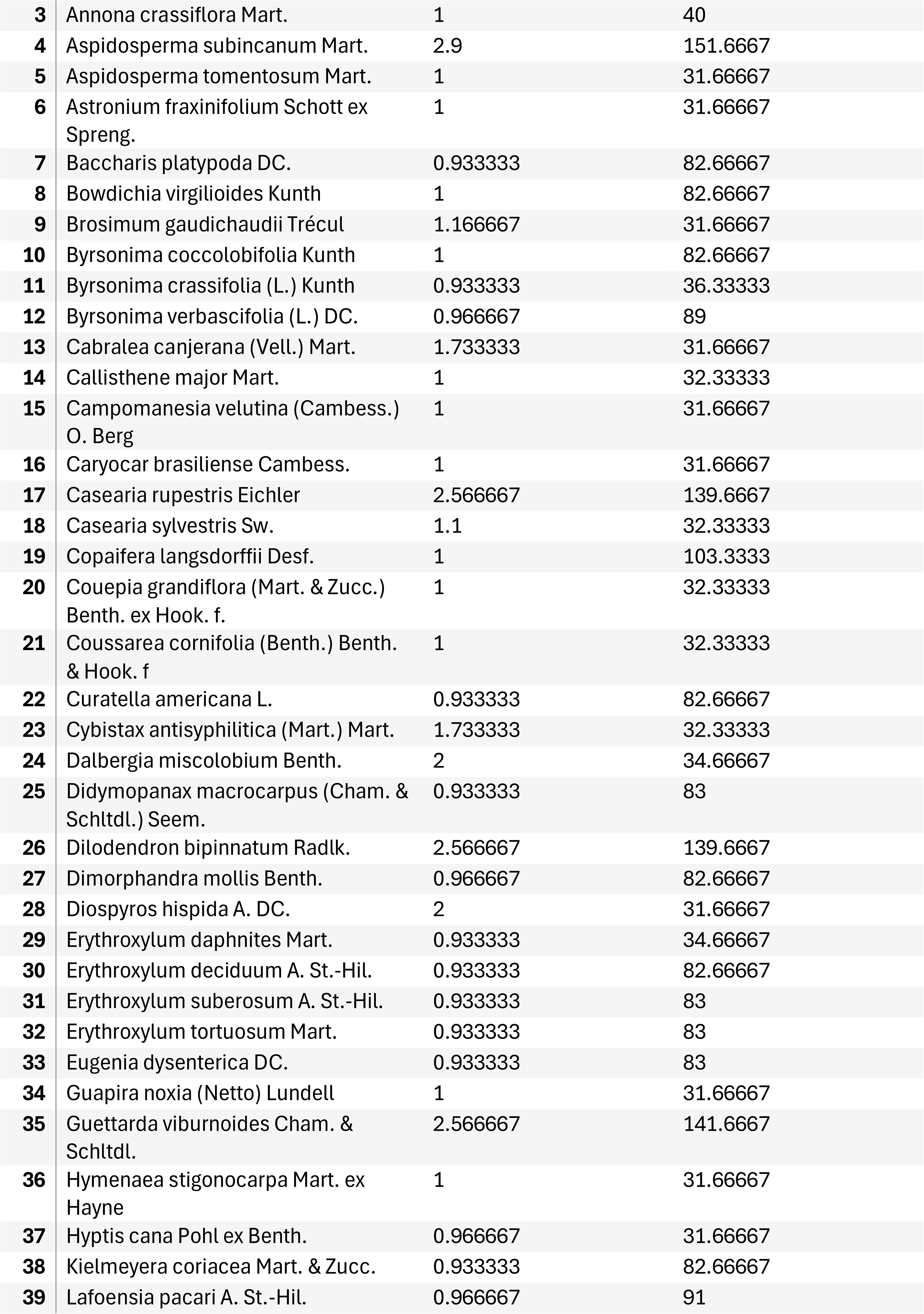
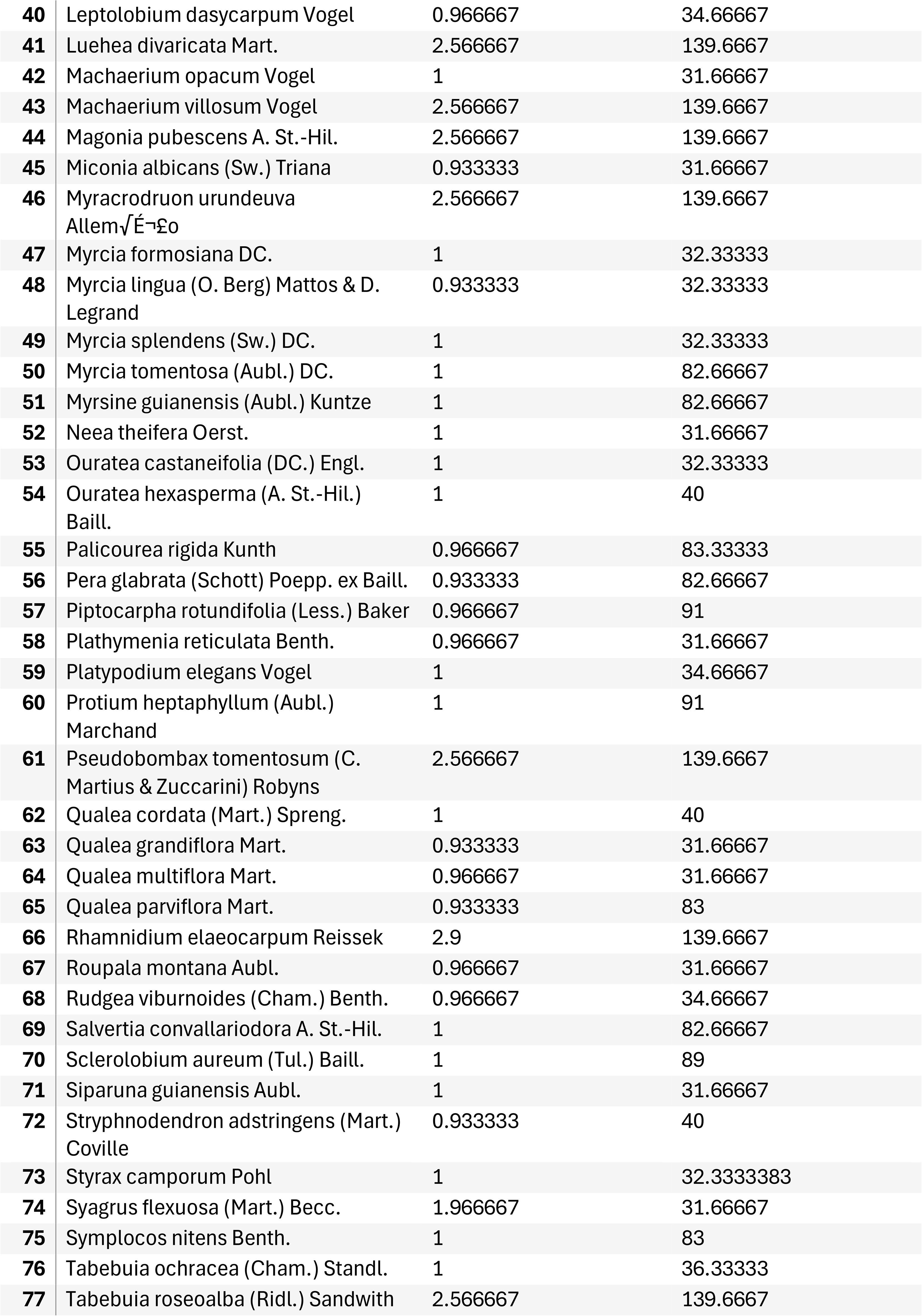
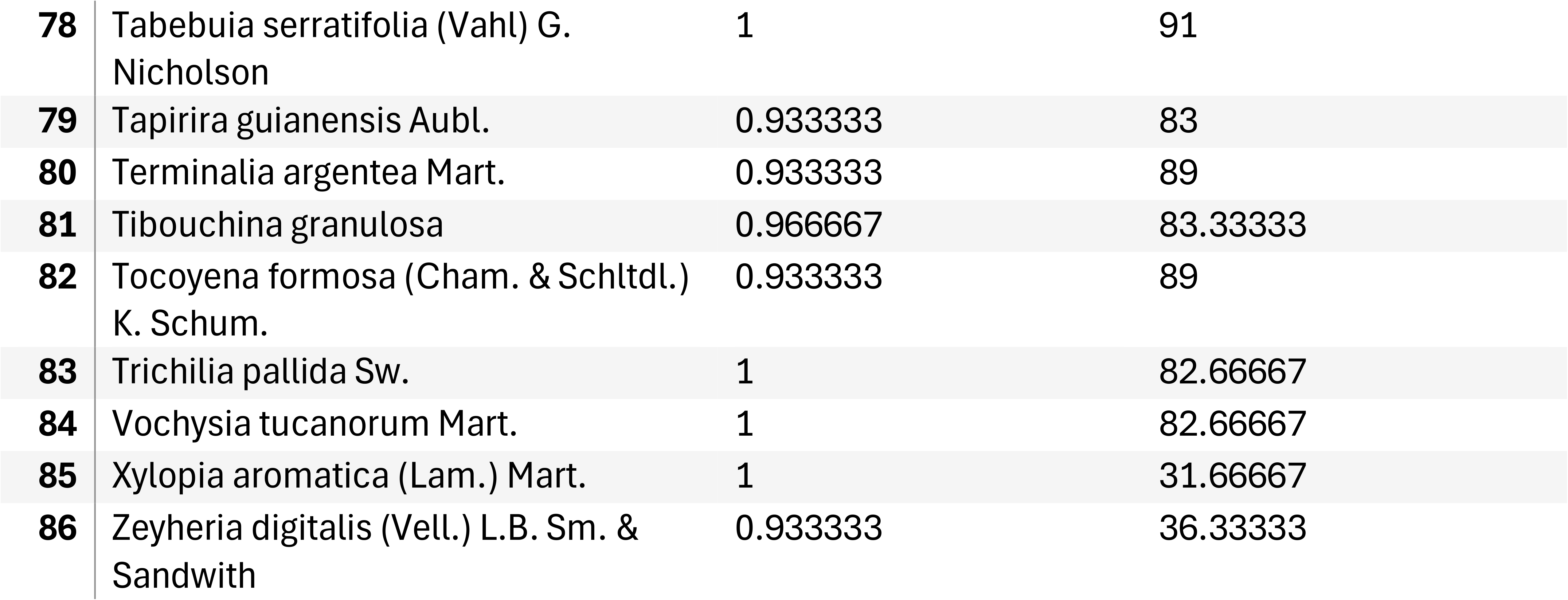
Species references and x, y coordinates of species in the Patchwork Biplot of Figures S2 and S3 of the case study. Only species with sampled individuals in at least three communities were considered for isoclines.

**Table S2.**
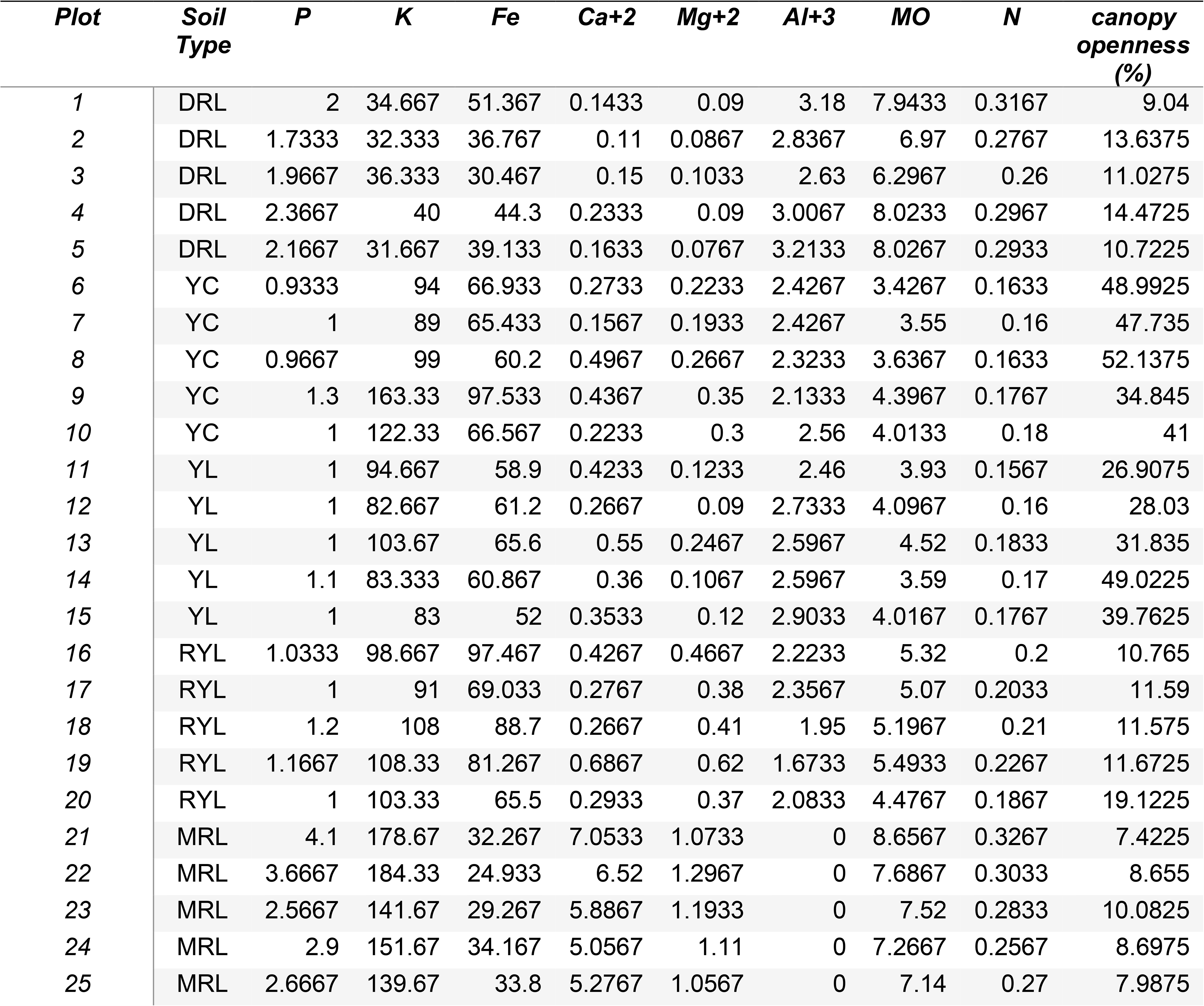
Plots, soil properties and canopy openness in the Paraopeba Reserve Cerrado. Soil contents (dag.kg^-1^): Dystrophic Red Latosol - DRL, Yellow Cambisol - YC, Yellow Latosol - YL, Red-Yellow Latosol - RYL, Mesotrophic Red Latosol - MRL.

