## Supplementary material for "A Multiscale Translation of Tilman’s R* and Its Empirical Application in the Cerrado"

1-LEEP - Laboratory of Ecology and Evolution of Plants, Universidade Federal de Viçosa (UFV), Viçosa, 36570-900
2-Botany Graduate Program, DBV, Universidade Federal de Viçosa, Viçosa, 36570-900
3-Ecology Graduate Program, CCB, Universidade Federal de Viçosa, Viçosa, 36570-900

Supplementary Material

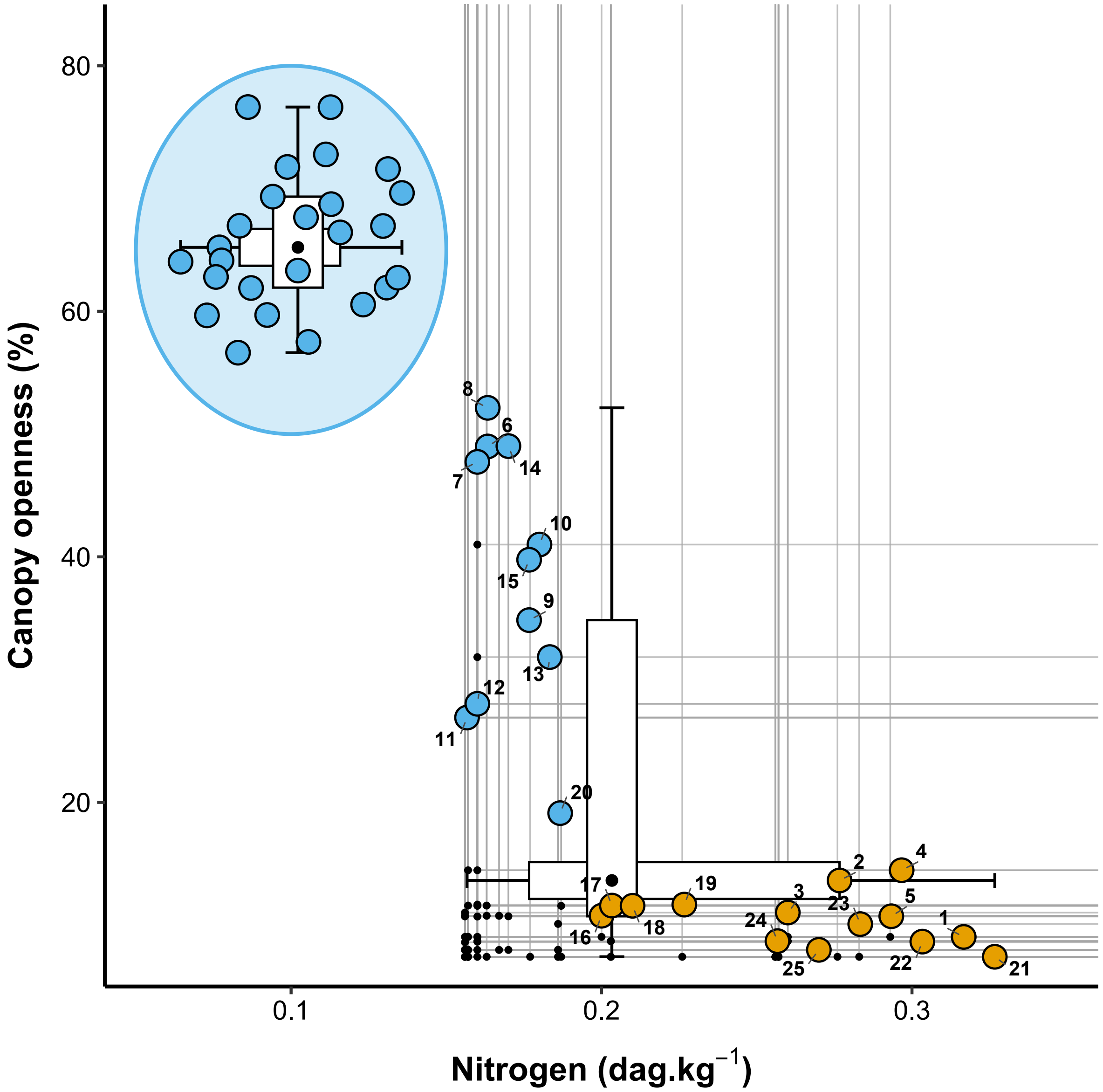


Figure S1 - Patchwork Biplot of plots in the Paraopeba Reserve Cerrado. Inside the blue ellipse: Simulation of the communities of the Paroapeba Reserve assuming N values reported by Belmok et al. (2019) for Cerrados with a fire regime in recent decades, and canopy openness values by (Ribeiro & Walter, 1998) for the expected range of variation for the Cerrado in general. Outside the blue ellipse: in the upper left quadrant of the Patchwork Biplot, the communities may still retain some past effect of disturbances prior to the protection of the Paroapeba Reserve, therefore in this figure they are in blue, indicating that they are not yet predominantly governed by stress; In the lower right quadrant, communities are organized along canopy openness contours, suggesting sliding along the contours as predicted in Figure 8. Past disturbances in regimes similar to those of the IBGE Reserve (large blue circle) increase canopy openness, increasing illumination at ground level and decreasing shading stresses. Communities in orange are under light limitation and governed by stress, in the lower rare quadrant. As the canopy openness, the values of N were obtained only for the Paraopeba Reserve; therefore, the communities in the blue ellipse did not define the isoclines.

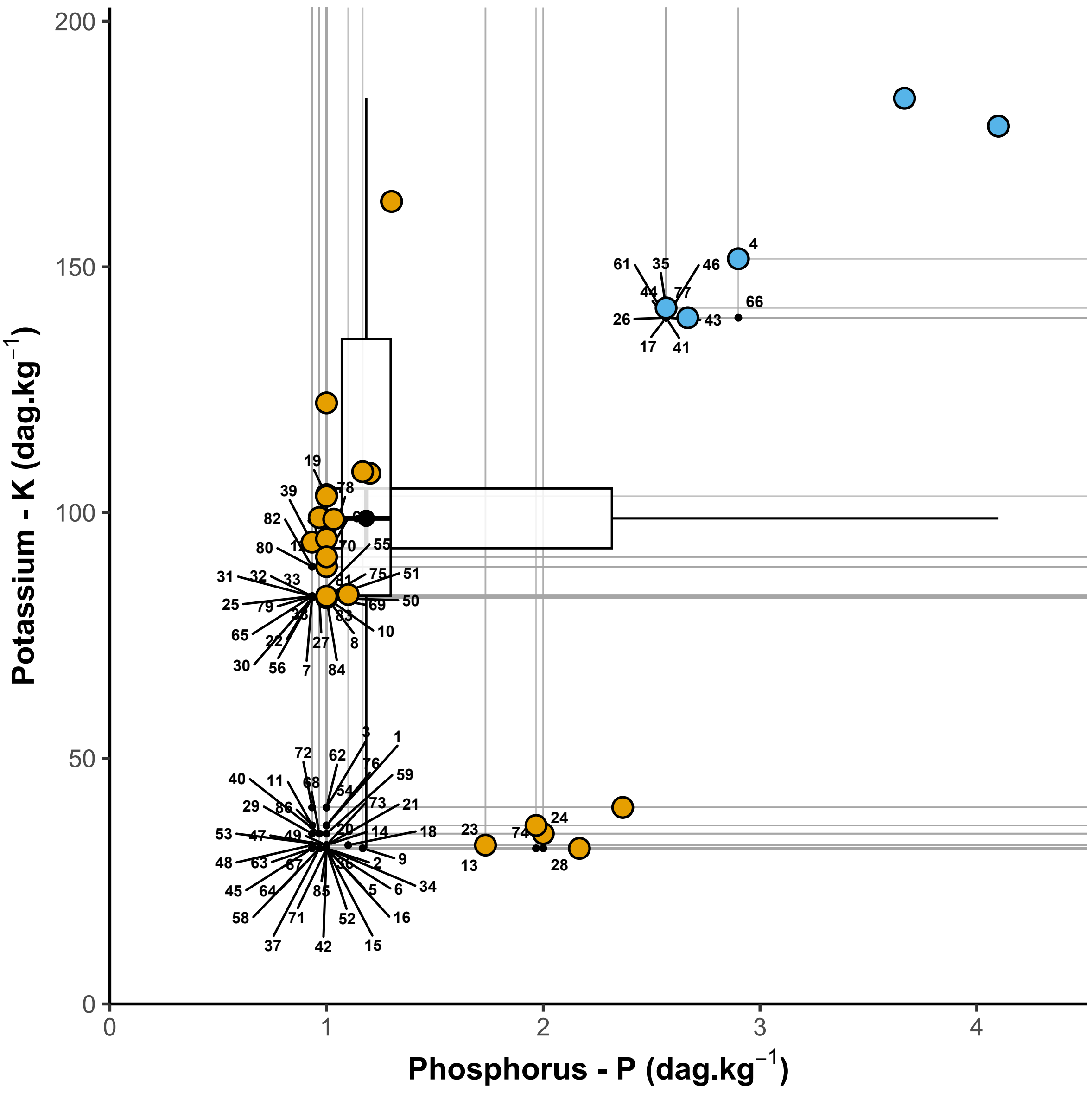


Figure S2 - Patchwork Biplot of plots in the Paraopeba Reserve Cerrado with numbers indicating species of the Table S1. In the upper right quadrant (Mainstream Quadrant) of the Patchwork Biplot, the communities may still retain some past effect of disturbances prior to the protection of the Paroapeba Reserve, therefore in this figure they are in blue, indicating that they are not yet predominantly governed by stress; In the lower right quadrant and in the left, communities are organized along phosphorus and potassium contours. Communities in orange are under phosphorus or potassium limitation and governed by stress.

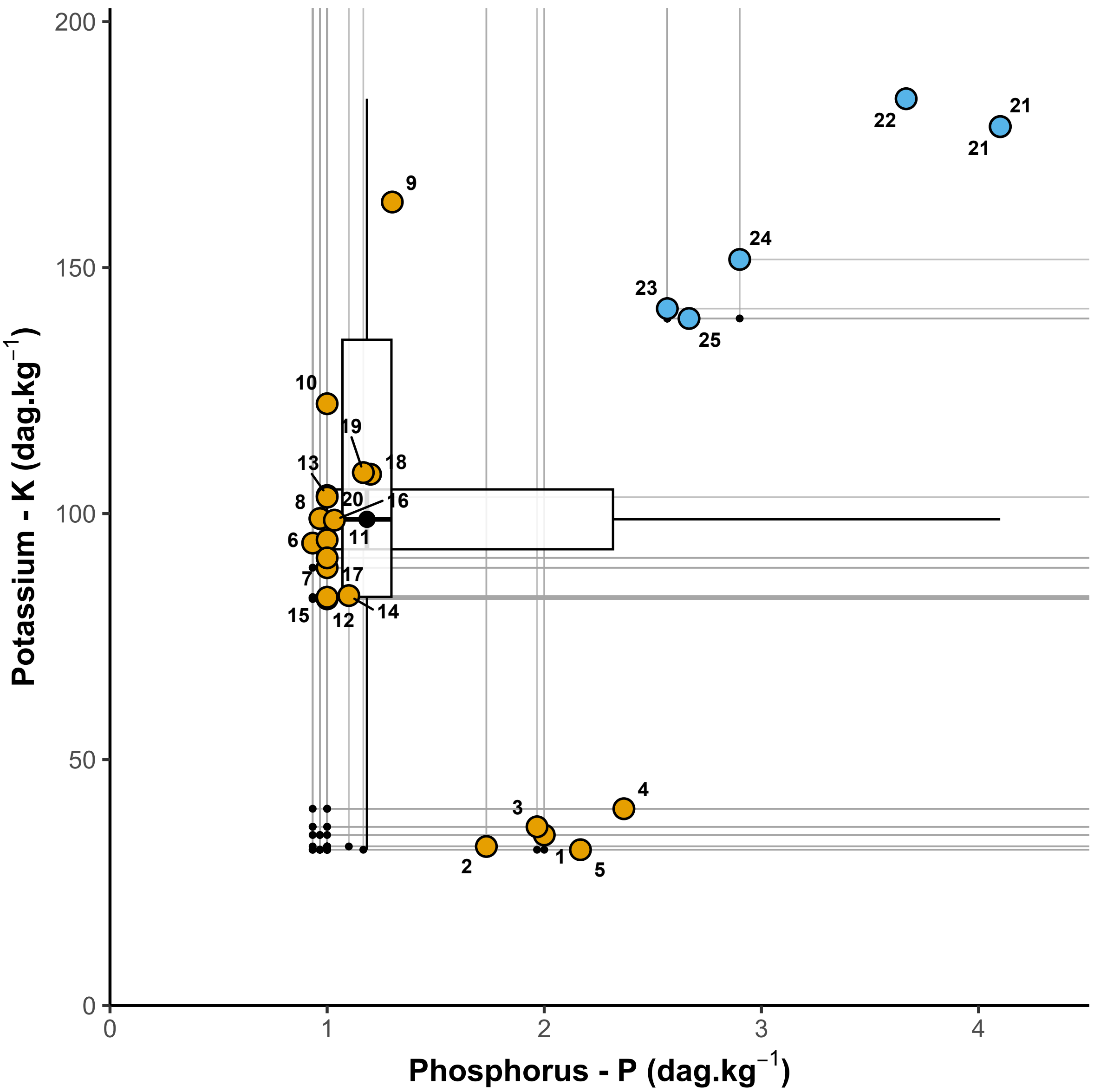


Figure S3 - Patchwork Biplot of plots in the Paraopeba Reserve Cerrado with numbers indicating the plots. In the upper right quadrant (Mainstream Quadrant) of the Patchwork Biplot, the communities may still retain some past effect of disturbances prior to the protection of the Paroapeba Reserve, therefore in this figure they are in blue, indicating that they are not yet predominantly governed by stress; In the lower right quadrant and in the left, communities are organized along phosphorus and potassium contours. Communities in orange are under phosphorus or potassium limitation and governed by stress.

Table S1 - Species references and x, y coordinates of species in the Patchwork Biplot of Figures S2 and S3 of the case study. Only species with sampled individuals in at least three communities were considered for isoclines

| # | Species | X axis (Phosphorus Dag.Kg$^{-1}$) | Y axis (Potassium Dag.Kg$^{-1}$) |
|---|---|---|---|
| 1 | Acacia polyphylla DC. | 1 | 36.33333 |
| 2 | Alibertia edulis (Rich.) A. Rich. ex DC. | 1 | 31.66667 |
| 3 | Annona crassiflora Mart. | 1 | 40 |
| 4 | Aspidosperma subincanum Mart. | 2.9 | 151.6667 |
| 5 | Aspidosperma tomentosum Mart. | 1 | 31.66667 |
| 6 | Astronium fraxinifolium Schott ex Spreng. | 1 | 31.66667 |
| 7 | Baccharis platypoda DC. | 0.933333 | 82.66667 |
| 8 | Bowdichia virgilioides Kunth | 1 | 82.66667 |
| 9 | Brosimum gaudichaudii Trécul | 1.166667 | 31.66667 |
| 10 | Byrsonima coccolobifolia Kunth | 1 | 82.66667 |
| 11 | Byrsonima crassifolia (L.) Kunth | 0.933333 | 36.33333 |
| 12 | Byrsonima verbascifolia (L.) DC. | 0.966667 | 89 |
| 13 | Cabralea canjerana (Vell.) Mart. | 1.733333 | 31.66667 |
| 14 | Callisthene major Mart. | 1 | 32.33333 |
| 15 | Campomanesia velutina (Cambess.) O. Berg | 1 | 31.66667 |
| 16 | Caryocar brasiliense Cambess. | 1 | 31.66667 |
| 17 | Casearia rupestris Eichler | 2.566667 | 139.6667 |
| 18 | Casearia sylvestris Sw. | 1.1 | 32.33333 |
| 19 | Copaifera langsdorffii Desf. | 1 | 103.3333 |
| 20 | Couepia grandiflora (Mart. & Zucc.) Benth. ex Hook. f. | 1 | 32.33333 |
| 21 | Coussarea cornifolia (Benth.) Benth. & Hook. f | 1 | 32.33333 |
| 22 | Curatella americana L. | 0.933333 | 82.66667 |
| 23 | Cybistax antisyphilitica (Mart.) Mart. | 1.733333 | 32.33333 |
| 24 | Dalbergia miscolobium Benth. | 2 | 34.66667 |
| 25 | Didymopanax macrocarpus (Cham. & Schltdl.) Seem. | 0.933333 | 83 |
| 26 | Dilodendron bipinnatum Radlk. | 2.566667 | 139.6667 |
| 27 | Dimorphandra mollis Benth. | 0.966667 | 82.66667 |
| 28 | Diospyros hispida A. DC. | 2 | 31.66667 |
| 29 | Erythroxylum daphnites Mart. | 0.933333 | 34.66667 |
| 30 | Erythroxylum deciduum A. St.-Hil. | 0.933333 | 82.66667 |
| 31 | Erythroxylum suberosum A. St.-Hil. | 0.933333 | 83 |
| 32 | Erythroxylum tortuosum Mart. | 0.933333 | 83 |
| 33 | Eugenia dysenterica DC. | 0.933333 | 83 |
| 34 | Guapira noxia (Netto) Lundell | 1 | 31.66667 |

| | | | |
|---|---|---|---|
| **35** | Guettarda viburnoides Cham. & Schltdl. | 2.566667 | 141.6667 |
| **36** | Hymenaea stigonocarpa Mart. ex Hayne | 1 | 31.66667 |
| **37** | Hyptis cana Pohl ex Benth. | 0.966667 | 31.66667 |
| **38** | Kielmeyera coriacea Mart. & Zucc. | 0.933333 | 82.66667 |
| **39** | Lafoensia pacari A. St.-Hil. | 0.966667 | 91 |
| **40** | Leptolobium dasycarpum Vogel | 0.966667 | 34.66667 |
| **41** | Luehea divaricata Mart. | 2.566667 | 139.6667 |
| **42** | Machaerium opacum Vogel | 1 | 31.66667 |
| **43** | Machaerium villosum Vogel | 2.566667 | 139.6667 |
| **44** | Magonia pubescens A. St.-Hil. | 2.566667 | 139.6667 |
| **45** | Miconia albicans (Sw.) Triana | 0.933333 | 31.66667 |
| **46** | Myracrodruon urundeuva Allem√É¬£o | 2.566667 | 139.6667 |
| **47** | Myrcia formosiana DC. | 1 | 32.33333 |
| **48** | Myrcia lingua (O. Berg) Mattos & D. Legrand | 0.933333 | 32.33333 |
| **49** | Myrcia splendens (Sw.) DC. | 1 | 32.33333 |
| **50** | Myrcia tomentosa (Aubl.) DC. | 1 | 82.66667 |
| **51** | Myrsine guianensis (Aubl.) Kuntze | 1 | 82.66667 |
| **52** | Neea theifera Oerst. | 1 | 31.66667 |
| **53** | Ouratea castaneifolia (DC.) Engl. | 1 | 32.33333 |
| **54** | Ouratea hexasperma (A. St.-Hil.) Baill. | 1 | 40 |
| **55** | Palicourea rigida Kunth | 0.966667 | 83.33333 |
| **56** | Pera glabrata (Schott) Poepp. ex Baill. | 0.933333 | 82.66667 |
| **57** | Piptocarpha rotundifolia (Less.) Baker | 0.966667 | 91 |
| **58** | Plathymenia reticulata Benth. | 0.966667 | 31.66667 |
| **59** | Platypodium elegans Vogel | 1 | 34.66667 |
| **60** | Protium heptaphyllum (Aubl.) Marchand | 1 | 91 |
| **61** | Pseudobombax tomentosum (C. Martius & Zuccarini) Robyns | 2.566667 | 139.6667 |
| **62** | Qualea cordata (Mart.) Spreng. | 1 | 40 |
| **63** | Qualea grandiflora Mart. | 0.933333 | 31.66667 |
| **64** | Qualea multiflora Mart. | 0.966667 | 31.66667 |
| **65** | Qualea parviflora Mart. | 0.933333 | 83 |
| **66** | Rhamnidium elaeocarpum Reissek | 2.9 | 139.6667 |
| **67** | Roupala montana Aubl. | 0.966667 | 31.66667 |
| **68** | Rudgea viburnoides (Cham.) Benth. | 0.966667 | 34.66667 |
| **69** | Salvertia convallariodora A. St.-Hil. | 1 | 82.66667 |
| **70** | Sclerolobium aureum (Tul.) Baill. | 1 | 89 |
| **71** | Siparuna guianensis Aubl. | 1 | 31.66667 |

| | | | |
|---|---|---|---|
| **72** | Stryphnodendron adstringens (Mart.) Coville | 0.933333 | 40 |
| **73** | Styrax camporum Pohl | 1 | 32.3333383 |
| **74** | Syagrus flexuosa (Mart.) Becc. | 1.966667 | 31.66667 |
| **75** | Symplocos nitens Benth. | 1 | 83 |
| **76** | Tabebuia ochracea (Cham.) Standl. | 1 | 36.33333 |
| **77** | Tabebuia roseoalba (Ridl.) Sandwith | 2.566667 | 139.6667 |
| **78** | Tabebuia serratifolia (Vahl) G. Nicholson | 1 | 91 |
| **79** | Tapirira guianensis Aubl. | 0.933333 | 83 |
| **80** | Terminalia argentea Mart. | 0.933333 | 89 |
| **81** | Tibouchina granulosa | 0.966667 | 83.33333 |
| **82** | Tocoyena formosa (Cham. & Schltdl.) K. Schum. | 0.933333 | 89 |
| **83** | Trichilia pallida Sw. | 1 | 82.66667 |
| **84** | Vochysia tucanorum Mart. | 1 | 82.66667 |
| **85** | Xylopia aromatica (Lam.) Mart. | 1 | 31.66667 |
| **86** | Zeyheria digitalis (Vell.) L.B. Sm. & Sandwith | 0.933333 | 36.33333 |

Table S2 - Plots, soil properties and canopy openness in the Paraopeba Reserve Cerrado. Soil contents (dag.kg$^{-1}$): Dystrophic Red Latosol - DRL, Yellow Cambisol - YC, Yellow Latosol - YL, Red-Yellow Latosol - RYL, Mesotrophic Red Latosol - MRL

| *Plot* | *Soil Type* | *P* | *K* | *Fe* | *Ca+2* | *Mg+2* | *Al+3* | *MO* | *N* | *canopy openness (%)* |
|---|---|---|---|---|---|---|---|---|---|---|
| *1* | DRL | 2 | 34.667 | 51.367 | 0.1433 | 0.09 | 3.18 | 7.9433 | 0.3167 | 9.04 |
| *2* | DRL | 1.7333 | 32.333 | 36.767 | 0.11 | 0.0867 | 2.8367 | 6.97 | 0.2767 | 13.6375 |
| *3* | DRL | 1.9667 | 36.333 | 30.467 | 0.15 | 0.1033 | 2.63 | 6.2967 | 0.26 | 11.0275 |
| *4* | DRL | 2.3667 | 40 | 44.3 | 0.2333 | 0.09 | 3.0067 | 8.0233 | 0.2967 | 14.4725 |
| *5* | DRL | 2.1667 | 31.667 | 39.133 | 0.1633 | 0.0767 | 3.2133 | 8.0267 | 0.2933 | 10.7225 |
| *6* | YC | 0.9333 | 94 | 66.933 | 0.2733 | 0.2233 | 2.4267 | 3.4267 | 0.1633 | 48.9925 |
| *7* | YC | 1 | 89 | 65.433 | 0.1567 | 0.1933 | 2.4267 | 3.55 | 0.16 | 47.735 |
| *8* | YC | 0.9667 | 99 | 60.2 | 0.4967 | 0.2667 | 2.3233 | 3.6367 | 0.1633 | 52.1375 |
| *9* | YC | 1.3 | 163.33 | 97.533 | 0.4367 | 0.35 | 2.1333 | 4.3967 | 0.1767 | 34.845 |
| *10* | YC | 1 | 122.33 | 66.567 | 0.2233 | 0.3 | 2.56 | 4.0133 | 0.18 | 41 |
| *11* | YL | 1 | 94.667 | 58.9 | 0.4233 | 0.1233 | 2.46 | 3.93 | 0.1567 | 26.9075 |
| *12* | YL | 1 | 82.667 | 61.2 | 0.2667 | 0.09 | 2.7333 | 4.0967 | 0.16 | 28.03 |
| *13* | YL | 1 | 103.67 | 65.6 | 0.55 | 0.2467 | 2.5967 | 4.52 | 0.1833 | 31.835 |
| *14* | YL | 1.1 | 83.333 | 60.867 | 0.36 | 0.1067 | 2.5967 | 3.59 | 0.17 | 49.0225 |
| *15* | YL | 1 | 83 | 52 | 0.3533 | 0.12 | 2.9033 | 4.0167 | 0.1767 | 39.7625 |
| *16* | RYL | 1.0333 | 98.667 | 97.467 | 0.4267 | 0.4667 | 2.2233 | 5.32 | 0.2 | 10.765 |
| *17* | RYL | 1 | 91 | 69.033 | 0.2767 | 0.38 | 2.3567 | 5.07 | 0.2033 | 11.59 |
| *18* | RYL | 1.2 | 108 | 88.7 | 0.2667 | 0.41 | 1.95 | 5.1967 | 0.21 | 11.575 |
| *19* | RYL | 1.1667 | 108.33 | 81.267 | 0.6867 | 0.62 | 1.6733 | 5.4933 | 0.2267 | 11.6725 |
| *20* | RYL | 1 | 103.33 | 65.5 | 0.2933 | 0.37 | 2.0833 | 4.4767 | 0.1867 | 19.1225 |
| *21* | MRL | 4.1 | 178.67 | 32.267 | 7.0533 | 1.0733 | 0 | 8.6567 | 0.3267 | 7.4225 |
| *22* | MRL | 3.6667 | 184.33 | 24.933 | 6.52 | 1.2967 | 0 | 7.6867 | 0.3033 | 8.655 |
| *23* | MRL | 2.5667 | 141.67 | 29.267 | 5.8867 | 1.1933 | 0 | 7.52 | 0.2833 | 10.0825 |
| *24* | MRL | 2.9 | 151.67 | 34.167 | 5.0567 | 1.11 | 0 | 7.2667 | 0.2567 | 8.6975 |
| *25* | MRL | 2.6667 | 139.67 | 33.8 | 5.2767 | 1.0567 | 0 | 7.14 | 0.27 | 7.9875 |
